# Combinatorial Optimization of Antibody Libraries via Constrained Integer Programming

**DOI:** 10.1101/2024.11.03.621763

**Authors:** Conor F. Hayes, Andre R. Goncalves, Steven Magana-Zook, Jacob Pettit, Ahmet Can Solak, Daniel Faissol, Mikel Landajuela

## Abstract

Designing effective antibody libraries is a challenging combinatorial search problem in computational biology. We propose a novel integer linear programming (ILP) method that explicitly controls diversity and affinity objectives when generating candidate libraries. Our approach formulates library design as a constrained optimization problem, where diversity parameters and predicted binding scores are encoded as ILP constraints and objectives. Predicted binding scores are obtained via *AI-guided mutational fitness profiling*, which combines protein language models and inverse folding tools to evaluate mutational effects. We demonstrate the method on cold-start design tasks for Trastuzumab, D44.1, and Spesolimab, showing that our optimized libraries outperform baseline designs in both predicted affinity and sequence diversity. This hybrid search-and-learning framework illustrates how constrained optimization and predictive modeling can be combined to deliver interpretable, high-quality solutions to antibody library engineering. Code is available at https://github.com/llnl/protlib-designer.

**ACM Reference Format:** Conor F. Hayes, Andre R. Goncalves, Steven Magana-Zook, Jacob Pettit, Ahmet Can Solak, Daniel Faissol, and Mikel Landajuela. 2026. Combinatorial Optimization of Antibody Libraries via Constrained Integer Programming. In *Proc. of the 25th International Conference on Autonomous Agents and Multiagent Systems (AAMAS 2026), Paphos, Cyprus, May 25 – 29, 2026*, IFAAMAS, 22 pages.

## 1 INTRODUCTION

Antibody-based therapeutics have revolutionized the treatment of a wide range of diseases, including cancer, autoimmune disorders, and infectious diseases [5]. To develop these drugs, researchers often rely on directed evolution, a process that involves experimentally screening large libraries of antibodies to identify candidates with desirable properties [33]. In the early stages of directed evolution for antibody drug discovery, a key challenge is to design a diverse library of potential antibodies for further experimental screening [36, 38]. This challenge involves identifying promising leads that exhibit high affinity to target antigens and possess favorable developability characteristics [15, 48]. Effective library design is also central to many Bayesian optimization methods used in antibody discovery [8, 36], a problem known as batch active learning [6].

Traditionally, the initial step to bootstrap the directed evolution process has relied on random mutagenesis, often informed by deep mutational scanning data [28, 34, 52, 53], or through biomolecular simulations of binding free energies [4, 8]. These methods have been successfully applied to generate libraries that are enriched with high-affinity antibodies [44, 52]. However, these methods required a large amount of experimental data and/or computational resources, which can be prohibitively costly and time-consuming. Additionally, these approaches often overlook the need to maintain diversity within the antibody library, which is crucial for exploring the vast sequence space and overcoming potential failure modes or systematic biases in prediction tools [3, 9, 56].

Recent advancements in deep learning applied to biological sequences [19, 35, 40] and structures [7, 23], or a combination of both [54], have shown great promise as in silico screening tools for antibody drug discovery. These methods leverage the power of machine learning to learn from evolutionary scale data and predict the effects of mutations on antibody properties, such as binding affinity, stability, and developability [22, 45].

In this paper, we propose a novel approach that combines recent advances in deep learning for protein engineering with integer linear programming (ILP) to design diverse and high-quality antibody libraries. We are interested in a *cold-start* setting (see Appendix A.3), where the objective is to design effective starting libraries without the need for experimental or computational fitness data. This setting is relevant for rapid response design scenarios against escape variants or new targets [8], where the availability of experimental data is limited or non-existent, and for seeding the directed evolution process with diverse and high-quality candidates [36]. We summarize our contributions as follows:

- We introduce a novel method for antibody library design that leverages constrained integer linear programming to generate high-quality libraries with explicit control over diversity parameters.
- We apply the method to the problem of cold-start antibody library design using in silico deep mutational scanning data from inverse folding and protein language models.
- We evaluate the performance of our approach on three case studies involving the design of antibody libraries for the Trastuzumab, D44.1 and Spesolimab antibodies, showing that our method outperforms existing methods in terms of library quality and diversity.

## 2 RELATED WORK

Below we discuss relevant work related to antibody library design.

### Antibody library design

Typically when designing antibody libraries, experimental deep mutational scanning data or simulation data is utilized. For example, in [34] the authors start with experimental single-site deep mutational scanning to create a combinatorial library of antibody with respect to the Trastuzumab antibody. The library containing over 31, 000 designs is validated via in vitro experiments to determine binding quality. Additionally, in [8], an antibody that had previously lost potency due to mutations on the SARS-COV-2 antigen, is computationally redesigned to restore potency by producing an antibody library for experimental validation using a sequence generator informed by simulated data obtained after a large scale simulation campaign. In contrast, our proposed approach utilizes relatively inexpensive machine learning models to generate libraries in a cold-start setting without the need for experimental or computational fitness data.

### Antibody library design with diversity

Recently, a number of methods have been proposed to explicitly include diversity when generating antibody libraries. In [26], the authors use a quality-diversity optimization approach called MAP-elites [37] to produce a library of high performing and diverse antibodies. However, the approach in [26], struggles to generate libraries of a pre-defined size and requires an additional down-selection step to generate a final library.

In [55], a constrained Bayesian optimization approach is proposed to strike a balance between binding affinity and thermostability while maintaining sequence diversity. This method optimizes a latent space representation learned by a variational autoencoder [27] to produce a fit and diverse batch of designs. To ensure diversity, “black-box” constraints are utilized with an additional Levenstein distance constraint. In contrast, our ILP formulation optimizes directly on the additive objective values produced from the ML driven models, and enforces diversity directly in sequence-space with respect to the defined constraints.

In the larger context of protein design, [9, 56] recently proposed a differentiable generative approach to jointly optimize for the expected score and diversity of the generated library. By relaxing the discrete optimization problem to a continuous one, the authors are able to utilize gradient-based optimization methods to generate libraries. However, the diversity is represented by the entropy of the generator, which may not be directly related to the desired sequence diversity.

### Integer linear programming for antibody optimization

Antibody optimization has been formulated as integer linear programming in [44] and is used in an optimization pipeline to design a library with broadly binding and stable antibodies with respect to 180 divergent HIV viral strains. While the proposed approach is successful, the optimization pipeline requires extensive simulation data to train an ML binding predictor. The proposed ILP is formulated to optimize the respective binding scores of the trained ML predictor. We note the ML predictors may struggle to generalize to targets outside of the problem under consideration. Therefore, to apply this formulation to a new target viral strain, new data must be generated and the ML model retrained. Additionally, the ILP defined in [44] does not include diversity as a constraint during optimization.

### Machine learning for antibody engineering

Masked language models, trained on protein sequences, have been used to predict mutations in [19] and [35], where the scores produced by the models convey the predictive effects of mutations on protein function. For example, in [19], the authors use the likelihood of a mutation of a wild-type antibody under an ensemble of masked language models to select mutations on a number of antibodies. The models select evolutionarily plausible mutations without any context about the target antigen.

The majority of research to date has focused on using scores generated from masked language models without any context about the target antigen. However, the approach proposed in this work utilizes scores from antibody inverse folding models that can interpret provided antigen information. In [23], the trained inverse folding model achieves strong correlations when predicting antibody-antigen binding affinity. As machine learning methods improve, we assume the ability of these models to convey the predictive effects of mutations on protein function will also improve.

## 3 METHOD

The antibody reengineering problem starts with a wild-type antibody sequence w = (*w*_1_, …, *w*_*L*_ ) of length *L*, where each *w*_*i*_ takes a value in the set of *M* = 20 amino acids *A* = {*a*_1_, … *a*_*M*_ }. The wildtype antibody might present weak binding to an epitope g (showed in purple in Figure 1(a)). The antibody interface is a set of *N* ≤*L* position indices r = *r*{_1_, …, *r*_*N*_ }where 1 ≤*r*_*i*_ ≤*L* (gray positions in Figure 1(a)). Typically, residues at positions r are in contact with the antigen g or have been deemed important for binding. In the following, we use the notation ℐ_*A*_ (*a*) to denote the index of the amino acid *a* in the set *A*.

**Figure 1:**
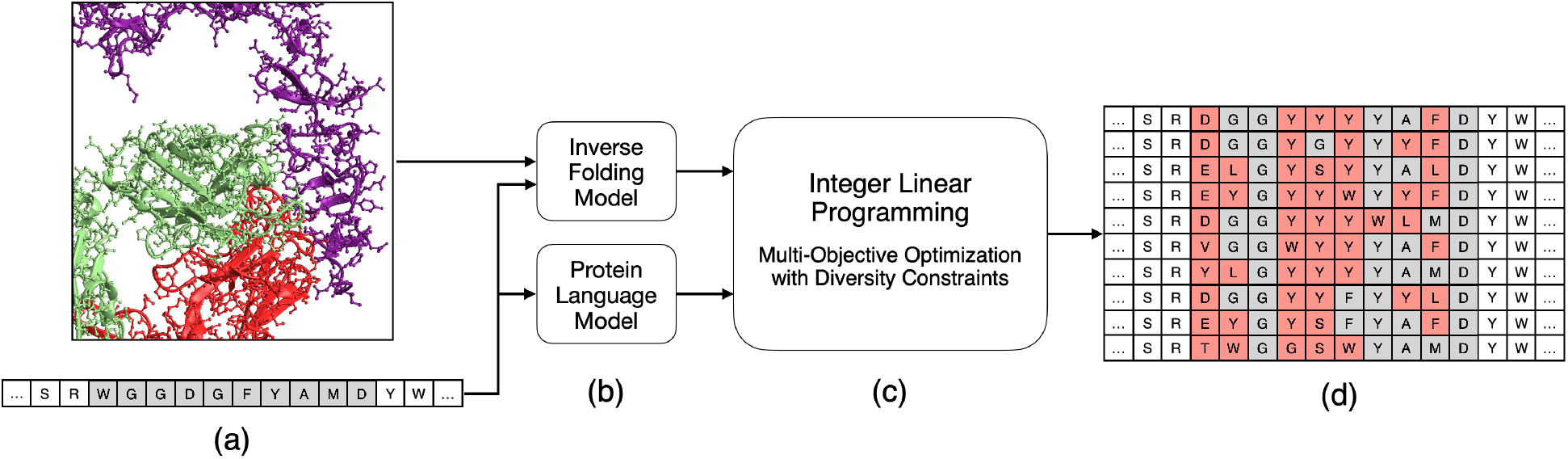
ProtLib-Designer: Schematic overview of the proposed method for antibody library design. (a) The input consists of an antibody-antigen complex and a target antibody sequence, defining the structural and sequence context for library generation. (b) In silico deep mutational scanning is performed using protein language models and inverse folding models to predict intrinsic and extrinsic mutational effects. (c) The generated mutational landscape is processed by a multi-objective linear programming solver, which balances multiple optimization criteria with diversity constraints. (d) The solver outputs an antibody library that is co-optimized for the predicted in silico scores while enforcing diversity constraints, ensuring broad sequence variation while maintaining functional relevance.

In a library design problem, the objective is to identify a set of mutants derived from the wild-type antibody w that exhibit enhanced binding to the antigen g while preserving developability properties [25, 42, 52]. Each mutant is represented by x = (*x*_1_, …, *x*_*N*_ ), where *x*_*i*_ ∈ *A*. Additionally, the set of mutants should be diverse to encompass a broad spectrum of potential antibodies and reduce the risk of experimental failure [49]. Consider a batch of *K* mutants *B* = x_1_, …, x_*K*_ . Formally, we can write the problem as the following constrained multi-objective optimization problem:

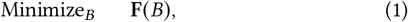

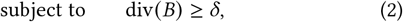

where F (*B*) is a vector of aggregated scores for the batch *B*, div *B* is the diversity of the batch *B*, and δ is a given tolerance. Different scoring functions F (*B*) with different fidelity levels can be used to evaluate the mutants in the batch *B*. For instance, F (*B*) could involve experimental data, computational models, or a combination of both. In this work, we focus on scoring functions based on deep learning models.

### 3.1 In Silico Deep Mutational Scanning with Deep Learning

Experimental deep mutational scanning (DMS) is a powerful technique to guide protein design by systematically mutating position indices r to all amino acid identities in the set *A* and measuring the effects of these mutations on protein function [28, 52, 53]. The result is a matrix of scores *s*_*ij*_, where *s*_*ij*_ is the effect of mutating the amino acid at position *i* to amino acid *j* . DMS has been successfully used to design libraries enriched with high-affinity antibodies [44, 52]. However, DMS can be prohibitively costly and time-consuming, and it may not be feasible for all proteins of interest. For that reason, we propose to use in silico DMS, obtained via deep learning methods, to predict the effects of mutations on protein properties. We use two recent types of deep learning models to predict the effects of mutations: sequence-based models and structure-based models ^1^.

#### Intrinsic fitness score from protein language models

Protein language models (PLM) [19, 35, 40], trained on evolutionary-scale datasets of protein sequences, can be used to capture the evolutionary rules that govern protein sequences. PLMs capture the *intrinsic fitness* of a protein sequence, which is related to its folding stability, thermostability, developability, and evolutionary plausibility [18, 19]. In this work, we use a protein masked language model [19] and compute the score as

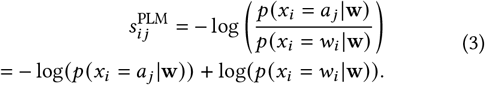

The score function (3) has been shown to be predictive of mutation effects [19, 35, 40]. In [35] they refer to (3) as the *wild-type marginal probability*, and they show extensive experiments on the predictive power of this score. Additionally, the computation of the predictive mutation effects for a given wildtype using the *wild-type marginal probability* only requires a single forward pass, making this approach computationally efficient and scalable as the number of positions increases. Note that (3) is a measure of the likelihood of observing a mutation to amino acid *a*_*j*_ at position *i* over the wild type sequence w. As such, it is formally related to the free energy difference between the wild type and the mutant state, 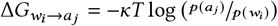, by the Boltzmann distribution in statistical mechanics, where *κ* is the Boltzmann constant and *T* is the temperature.

##### Algorithm 1

ProtLib-Designer: Diversity-Constrained Library Optimization

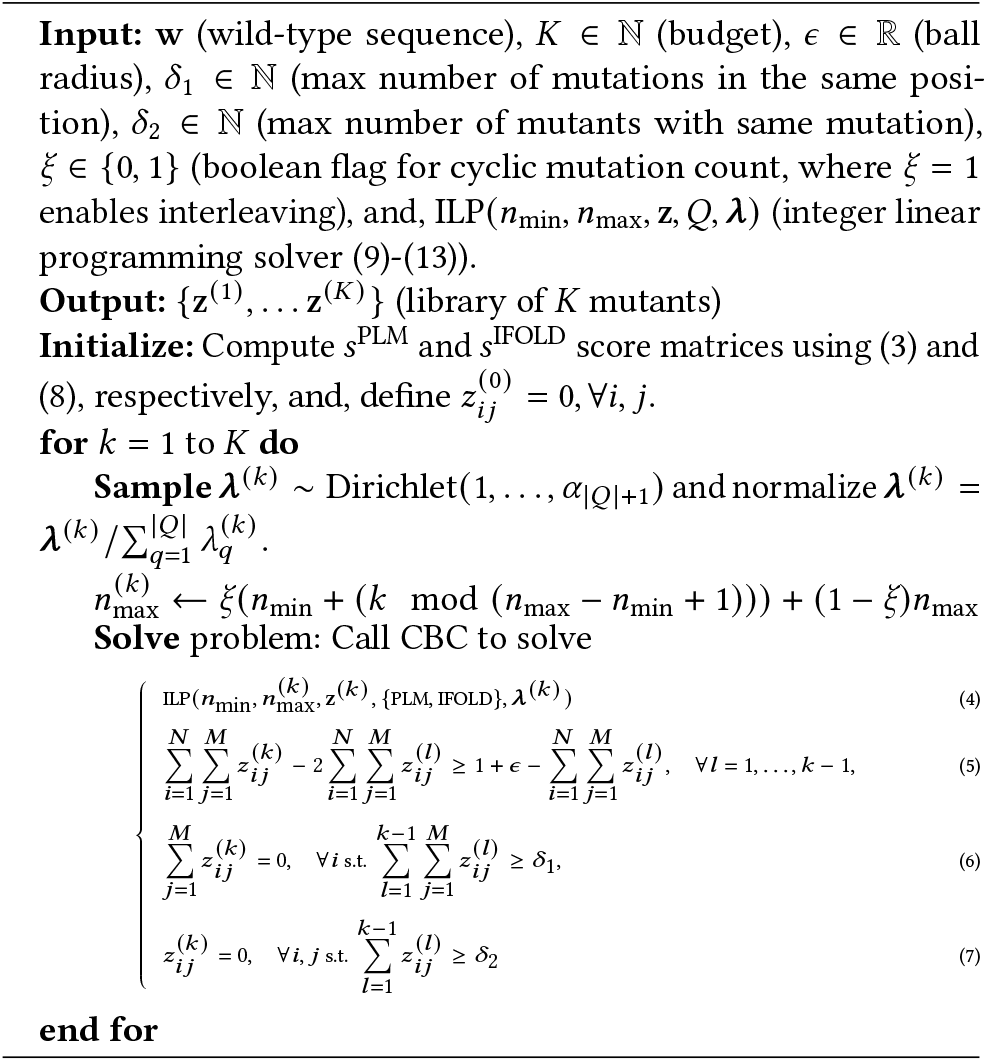

#### Extrinsic fitness score from inverse folding models

Several inversefolding (IFOLD) methods have been proposed to design protein sequences that fold into a given target structure [7, 21, 47]. It has been shown that, given an epitope as context, these methods can capture the *extrinsic fitness* of an antibody sequence, e.g., the specific selection pressure for binding to the epitope [18]. These models are conditioned on structural data of the co-complex structure struct(w, g). We define the score

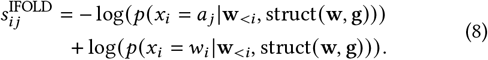

The score (8) has been investigated in [51] for a number of IFOLD methods. In this work, we consider Antifold [23], a method based on ESM-IF1 [21], fine-tuned on a large dataset of antibody data [30]. In the case of Antifold, struct w, g is given by the spatial coordinates of the backbone atoms (*N, C*_α_ and *C*). Geometric deep learning methods [22, 45] trained as regressors of antibody-antigen binding affinity could also be used to capture extrinsic fitness.

### 3.2 Multi-Objective Integer Linear Programming for Mutant Optimization

In this section, we consider the problem of finding a mutant of the wild-type antibody that minimizes a set *Q* of objectives. A mutant x = (*x*_1_, …, *x*_*N*_) is represented by a matrix z = *z*_*ij*_ ∈ {0, 1}, ∀1≤*i* ≤*N*, 1 ≤*j* ≤*M*, where *z*_*ij*_ = 1 if the amino acid *j* is present in the position *i* of the mutant, and *z*_*ij*_ = 0 otherwise. We assume that for each objective *q* ∈ *Q*, we have a score 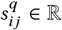 associated with the contribution of the amino acid *j* in position *i* to the objective *q*. We propose to frame the problem as a *multi-objective integer linear program* (ILP) as follows:

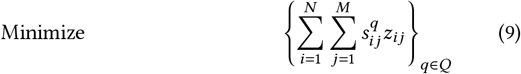

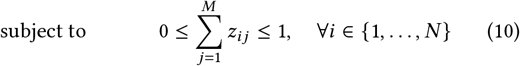

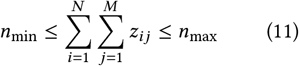

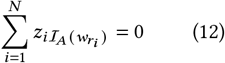

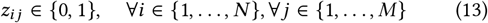

where (10) constraints the solution to one mutation per position, (11) constraints the number of mutations to be between *n*_min_ and *n*_max_, (12) constraints the current solution to be different from the wild-type at the positions r, and (13) constraints the solution to be binary.

If |*Q*| = 1, problem (9)–(13) can be solved by presenting it as an integer linear program. Specifically, problem (9)–(13) is a relaxed version of an assignment problem [29], where the bijectivity constraint is removed from (10). The convex optimization problem can be solved globally and efficiently using any available ILP solver. In this work, we use the COIN-OR Branch and Cut solver (CBC) [13]. If |*Q*| *>* 1, the problem is a multi-objective optimization problem that can be solved in a Pareto optimal sense using the weighted sum method [12], i.e., a single objective problem is solved by weighting the vector (9) with weights **λ** = (λ_1_, …, λ_|*Q* |_ ), where λ_*q*_ ≥ 0 and 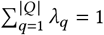. In the following, we consider *Q* = PLM, IFOLD with the corresponding score matrices introduced in (3) and (8). Note that for *q* = PLM, the additive model in (9) has been used in [35].

### 3.3 ProtLib-Designer: Diversity-Constrained Library Optimization

We now combine the ideas presented in the previous sections to propose a novel framework to solve the antibody library design problem with diversity constraints (1)–(2). We use the notation ILP (*n*_min_, *n*_max_, z, *Q*, λ) to represent the problem defined in equations (9)–(13) with minimum and maximum number of mutations per mutant given by *n*_min_ and *n*_max_, respectively, and objective weights λ. Given a budget of *K >* 0 mutants, we aim to find the top *K* solutions to the problem, ensuring diversity among the selected mutants.

We call the proposed algorithm *ProtLib-Designer* (PLD) and present it in Algorithm 1. PLD follows a *solve-and-remove* sequential strategy, where we *solve* a sequence of *K* problems (4), each supplemented with constraints (5)–(7) that depend on the solutions found in previous steps and that effectively *remove* areas of the search space (see Figure 3 for a step by step illustration). The *ball constraint* (5) ensures that a set of balls of radius ε around the solutions found in previous steps are removed from the feasible region of the next step. The *position constraint* (6) and the *mutation constraint* (7) ensure that the final antibody library presents at most δ_1_ mutations in the same position and at most δ_2_ mutants with the same mutation, respectively. Finally, the *cyclic mutation count* flag ξ allows for the cyclic variation of the maximum number of mutations per mutant, which can be useful to explore different levels of diversity in the library.

A key advantage of PLD is its minimal reliance on hyperparameters. The algorithm *requires only a small set of interpretable parameters*, including the ball radius ε and the diversity thresholds (δ_1_, δ_2_) (see the appendix). This simplicity enhances usability while maintaining flexibility, allowing researchers to tailor library design to specific experimental constraints without extensive hyperparameter tuning.

## 4 EXPERIMENTS

We demonstrate the flexibility of our library design approach by evaluating three antibody systems: Trastuzumab in complex with human epidermal growth factor receptor 2 (HER2) [34], the D44.1 antibody in complex with hen egg lysozyme (HEL) [34], and Spesolimab in complex with the IL36R extracellular domain receptor [46]. Due to space limitations, extensive ablations and baseline comparisons can be found in the appendix.

### Experimental Configuration

For evaluation, we compare the PLD algorithm with the *Linear Mutant Generator* (LMG) algorithm [8], the *Strength Pareto Evolutionary Algorithm 2* (SPEA2) [58], and the *ML-optimized library design with improved fitness and diversity* (MODIFY) method [9]. See the Appendix for extensive details on baselines, including implementation details. We also include the *multi-objective GFlowNet framework* [24] as an additional baseline in the appendix. We evaluated each algorithm on a series of metrics, including *expected utility of the batch* (BEU) and *hypervolume* (HV) (see the appendix for definitions). In addition, for the Trastuzumab system, we use a data-driven model to predict the binding affinity of each sequence to the target antigen (see the appendix for details).

For each antibody system we define a set of mutable positions—10 for Trastuzumab, 34 (17 H + 17 L) for D44.1, and 47 (H + L) for Spesolimab—and allow all 19 non–wild-type amino acids at each position (see the Appendix for full details). These search-space definitions drive the ILP diversity constraints in Eqs. (6)–(7), which bound per-position mutation counts and overall mutation frequencies to ensure both diversity and controlled mutation distributions.

For each system, we generate a batch *K* = 1, 000 mutated sequences from the wild-type. The number of mutations per sequence is constrained between *n*_min_ = 5 and *n*_max_ = 8. For all the baselines, we use objective (9) with Antifold (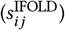) [23] and ProtBERT (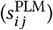) scores.

### 4.1 Results

In the following, we present the results of the PLD algorithm on the Trastuzumab, D44.1 and Spesolimab systems. In tables 1 to 3, the “Avg. rank” column reports the mean of each method’s 1-based ranks (1 = best)—computed across all the metrics after excluding the Pareto Front row—and serves as a single aggregated measure of overall library quality.

**Table 1:**
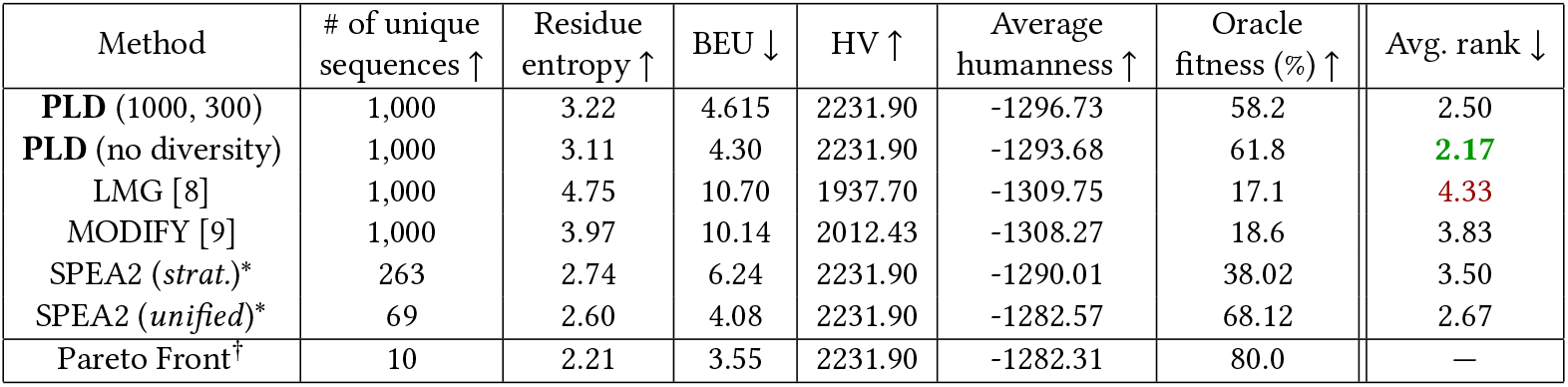
Diversity and fitness of libraries generated by each algorithm for Trastuzumab. ^∗^SPEA2 methods [58] were unable to generate libraries of the required size, making direct comparison with other algorithms difficult, though we include their results for reference. ^†^We approximate the Pareto front by combining the Pareto front of each algorithm’s solution set and removing dominated solutions.

| Method | # of unique sequences ↑ | Residue entropy ↑ | BEU ↓ | HV ↑ | Average humanness ↑ | Oracle fitness (%) ↑ | Avg. rank ↓ |
| --- | --- | --- | --- | --- | --- | --- | --- |
| PLD (1000, 300) | 1,000 | 3.22 | 4.615 | 2231.90 | -1296.73 | 58.2 | 2.50 |
| PLD (no diversity) | 1,000 | 3.11 | 4.30 | 2231.90 | -1293.68 | 61.8 | 2.17 |
| LMG [8] | 1,000 | 4.75 | 10.70 | 1937.70 | -1309.75 | 17.1 | 4.33 |
| MODIFY [9] | 1,000 | 3.97 | 10.14 | 2012.43 | -1308.27 | 18.6 | 3.83 |
| SPEA2 (strat.)* | 263 | 2.74 | 6.24 | 2231.90 | -1290.01 | 38.02 | 3.50 |
| SPEA2 (unified)* | 69 | 2.60 | 4.08 | 2231.90 | -1282.57 | 68.12 | 2.67 |
| Pareto Front<sup>†</sup> | 10 | 2.21 | 3.55 | 2231.90 | -1282.31 | 80.0 | — |

#### 4.1.1 Trastuzumab

We run PLD and baseline algorithms on the Trastuzumab system. In Figure 2, we show the libraries generated by each algorithm projected onto the objective space. Table 1 presents the compiled evaluation metrics.

**Figure 2:**
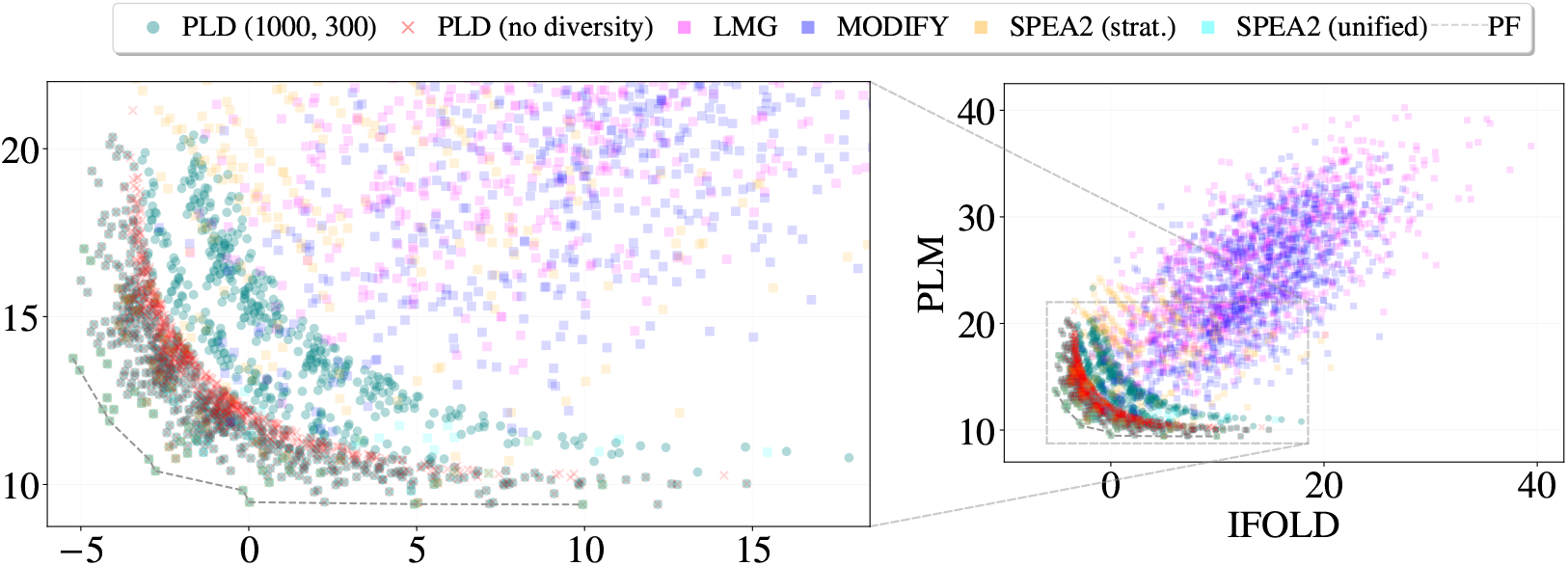
(right) Objective values of mutants generated by ProtLib-Designer (PLD) and baseline methods for Trastuzumab in complex with the HER2 receptor. Each point represents a 5 to 8-point mutant. (left) Zoomed-in perspective focusing on the objective values of the Pareto front and mutants generated by PLD with, and without, diversity constraints.

**Figure 3:**
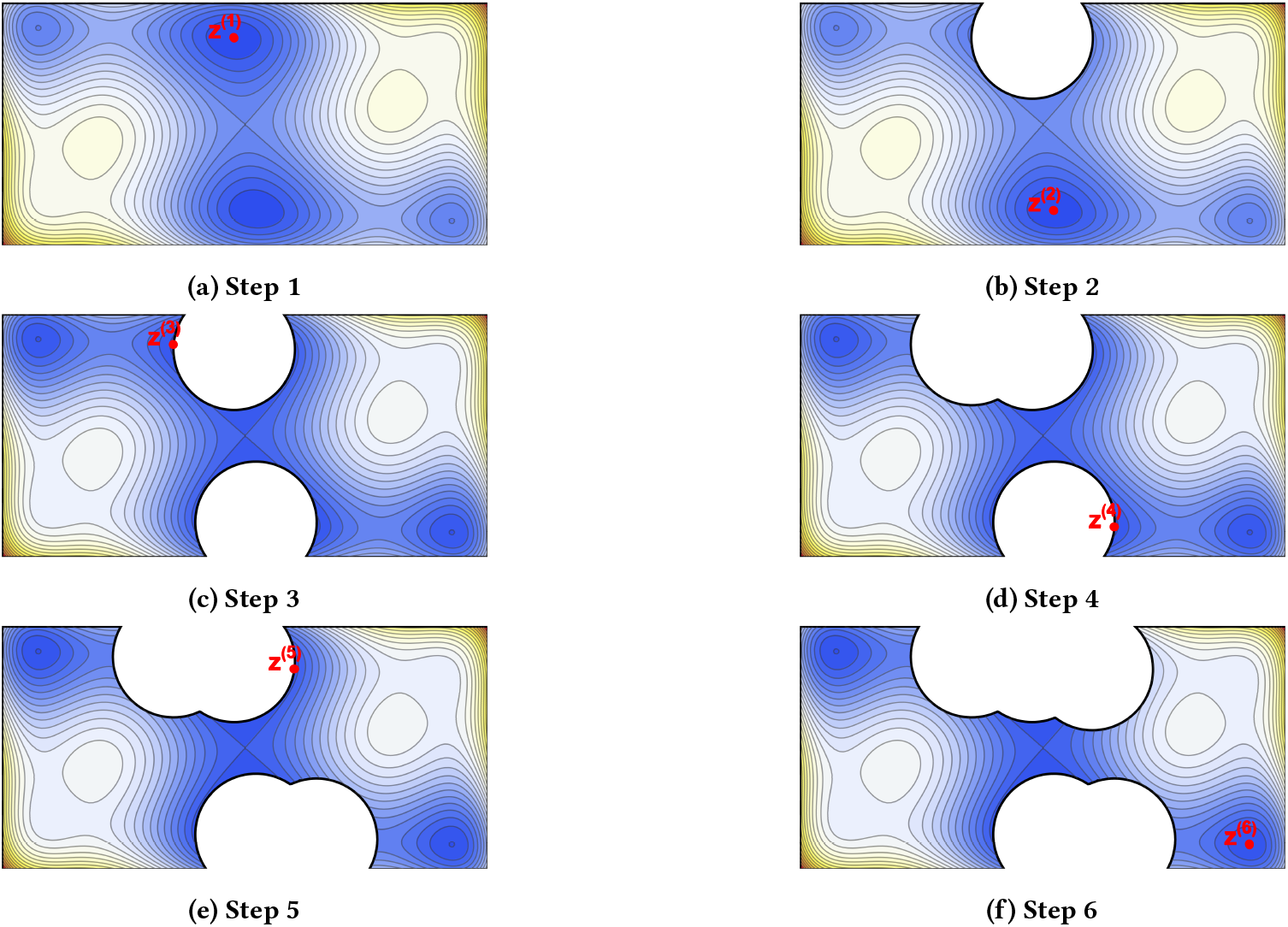
Illustration of the solve-and-remove strategy employed by ProtLib-Designer (PLD) to enforce diversity in antibody library design. The algorithm iteratively identifies the optimal solution within the current search space and removes a neighborhood of solutions around it (represented as a ball of radius ε) to prevent redundancy. By systematically eliminating similar solutions, PLD ensures that subsequent optimization steps explore new, diverse regions of the solution space. This process repeats until the desired number of unique solutions is obtained, balancing both optimization and diversity constraints.

In Figure 2 we observe that the PLD libraries are concentrated in the bottom-left corner of the objective space, near the Pareto front (PF). The mutants obtained by PLD with diversity constraints are organized in *stratified layers* separated by the effect of diversity constraints, while PLD without diversity constraints presents a single layer of mutants. In contrast, the LMG and MODIFY libraries distributions resemble the trace of a *shotgun* across the objective space. The SPEA2 libraries exhibit intermediate performance but face a significant limitation in generating the required library size, as shown in table 1. This is due to the presence of non-unique sequences in the final population (targeted at 1,000 individuals), resulting in less diverse libraries. For the problem of designing a 1, 000-mutant library, the PLD variants outperform all other methods in multi-objective metrics BEU and HV (see the appendix for definitions) while maintaining entropy values larger than 3. A tradeoff between entropy and BEU values is observed in the PLD libraries with and without diversity constraints.

Figure 4a and Table 1 compare Pareto fronts (HV relative to V_*ref*_ = [50, 50] ) for PLD and baselines. PLD and SPEA2 achieve the highest HV, span the widest objective ranges, and maintain diversity, outperforming all other methods. Moreover, both PLD variants obtained the best average rank scores, indicating superior overall library quality when aggregating across all metrics.

**Figure 4:**
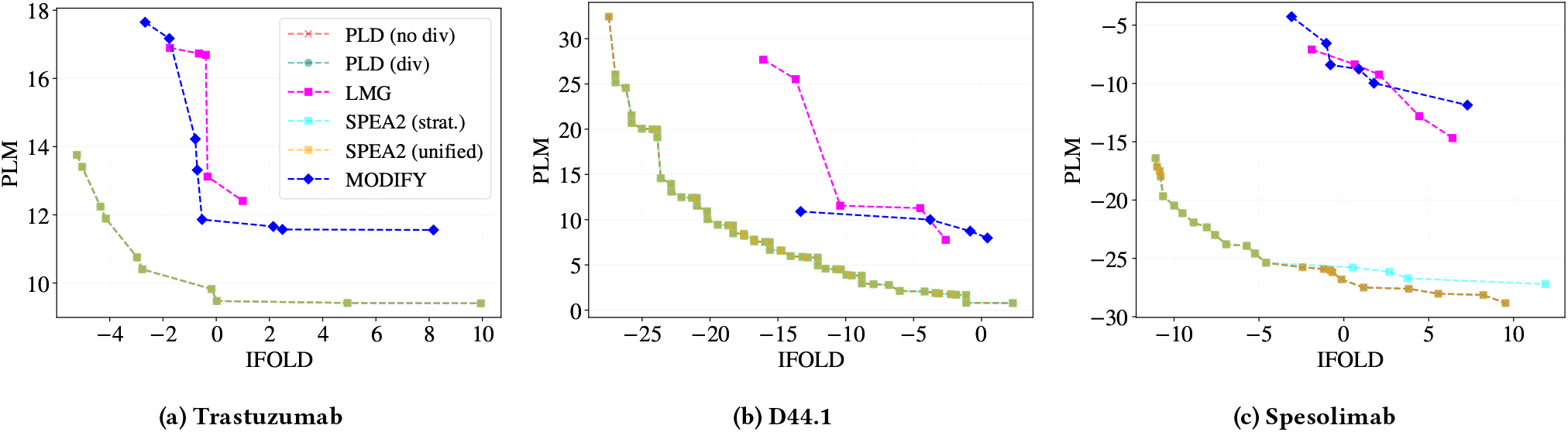
The individually computed Pareto front of the batches generated by ProtLib-Designer (PLD) and all baseline algorithms for Trastuzumab, D44.1, and Spesolimab. Note that the Pareto fronts obtained by PLD and SPEA2 (strat.) are identical in all systems.

The humanness and fitness columns in table 1 are used as external evaluation metrics for the library design problem. Following [16], we consider the log-likelihood of the mutants under ProtGPT2 as a proxy for developability (called average humanness). The oracle fitness is the percentage of mutants predicted to bind to the target antigen by a ML-classifier trained on experimental data (see the appendix for details). We observe ≃3.5-fold increase in predicted oracle fitness of the PLD mutants with higher average humanness compared to the LMG and MODIFY mutants. A similar trade-off between entropy and oracle fitness as described above is observed within the PLD libraries. The PLD libraries present a comparable predicted oracle fitness to the SPEA2 libraries, while involving a 5 to 14 times larger libraries. Note that the derived PF has an oracle fitness of 80%, justifying the choice of scoring functions. Overall, PLD provides the best libraries under the multi-objective metrics BEU and HV, oracle fitness, and average humanness, while providing library size control and flexibility to adjust the trade-off between diversity and fitness.

#### 4.1.2 D44.1

We evaluate PLD and baseline algorithms on the D44.1 system, following the same methodology as for Trastuzumab. Figure 4b shows the Pareto fronts obtained by each method, while Table 2 summarizes key performance metrics. Given the higher number of mutable positions in D44.1 (34) compared to Trastuzumab (10), the search space is significantly larger, resulting in a more complex and, in this case, a densely populated Pareto front. Despite this increased challenge, both PLD and SPEA2 effectively recovered the Pareto front, achieving good coverage of solutions, as can be seen in Figure 4b, highlighting the robustness and scalability of the proposed methodology.

**Table 2:** Comparison of diversity and fitness across libraries generated by each algorithm for D44.1. ^∗^SPEA2 methods [58] failed to produce libraries of the required size, complicating direct comparisons with other algorithms, though their results are included for reference.

| Method | # of unique sequences ↑ | Residue entropy ↑ | BEU ↓ | HV ↑ | Average humanness ↑ | Avg. rank ↓ |
| --- | --- | --- | --- | --- | --- | --- |
| **PLD** (1000, 500) | 1,000 | 2.93 | 1.84 | 1262.58 | -1815.90 | **2.0** |
| **PLD** (no diversity) | 1,000 | 2.81 | 1.41 | 1262.58 | -1821.62 | 2.8 |
| LMG [8] | 1,000 | 5.42 | 10.81 | 705.80 | -1830.79 | 3.6 |
| MODIFY [9] | 1,000 | 4.75 | 12.60 | 711.90 | -1838.80 | **4.0** |
| SPEA2 (*strat.*)* | 738 | 2.72 | 1.12 | 1262.58 | -1819.05 | 3.2 |
| SPEA2 (*unified*)* | 894 | 2.83 | 1.17 | 1258.93 | -1820.27 | 3.6 |
| Pareto Front<sup>†</sup> | 55 | 2.42 | 0.61 | 1262.58 | -1819.58 | — |

Similar to Trastuzumab, PLD libraries concentrate near the Pareto front, with diversity constraints forming stratified layers. SPEA2 struggles with library size, loosing up to 26% of the desired library size due to non-unique sequences. PLD achieves comparable BEU and HV scores to the Pareto front, while maintaining higher entropy values and average humanness scores, demonstrating its effectiveness in balancing diversity and optimization. Notably, PLD also achieves the best average rank, reflecting consistent overall performance across all metrics.

#### 4.1.3. Spesolimab

We evaluate PLD alongside baseline algorithms on the Spesolimab system [46], using the same methodology as above. Figure 4c displays each method’s Pareto front, and Table 3 reports the corresponding performance metrics. With 47 mutable positions—compared to 34 in D44.1 and 10 in Trastuzumab—the expanded search space produces a noticeably wider separation between the optimal front and those found by other baselines, a trend captured by the hypervolume values in Table 3. Importantly, only PLD and SPEA2 (stratified) fully recover the true Pareto front, highlighting our approach’s robustness and scalability in larger design problems.

**Table 3:** Comparison of diversity and fitness across libraries generated by each algorithm for Spesolimab. ^∗^SPEA2 (strat.) produced more than the required library size (1,810 instead of 1,000) by running four SPEA2 instances with different mutation counts (5–8) and then sampling down to 1,000 sequences (preserving Pareto-optimal ones). ^†^The Pareto front is approximated by merging all algorithms’ Pareto fronts and filtering out dominated solutions.

| Method | # of unique sequences ↑ | Residue entropy ↑ | BEU ↓ | HV ↑ | Average humanness ↑ | Avg. rank ↓ |
| --- | --- | --- | --- | --- | --- | --- |
| **PLD** (1000, 500) | 1,000 | 3.03 | -7.28 | 552.33 | -940.26 | **1.8** |
| **PLD** (no diversity) | 1,000 | 2.97 | -7.89 | 552.33 | -945.05 | 2.6 |
| LMG [8] | 1,000 | 6.01 | 4.50 | 118.72 | -949.41 | **3.8** |
| MODIFY [9] | 1,000 | 5.68 | 2.24 | 89.79 | -948.63 | **3.8** |
| SPEA2 (*strat.*)* | 1,000 | 2.99 | -6.51 | 552.33 | -942.55 | 2.6 |
| SPEA2 (*unified*)* | 1,000 | 3.10 | -6.19 | 528.02 | -941.55 | 2.8 |
| Pareto Front<sup>†</sup> | 24 | 2.65 | -8.53 | 552.33 | -945.53 | — |

## 5 CONCLUSION AND FUTURE WORK

In this work, we proposed ProtLib-Designer, a novel method for antibody library design that combines multi-objective optimization with diversity constraints. ProtLib-Designer leverages recent advances in sequence and structure-based machine learning models to compute in silico deep mutational scanning data. This data is then fed into an integer linear programming solver to generate diverse and high-performing antibody libraries. In an extensive evaluation involving three different antibodies, we showed that our method provides the best libraries in terms of multi-objective metrics, oracle fitness, and average humanness, while providing library size control and diversity-fitness trade-off flexibility.

As limitations, it is important to note that our method requires the structure of the antibody-antigen complex (which may not always be available) and that the quality of the generated libraries is affected by the quality of the scores predicted by the deep learning models. Also, the method might become computationally expensive for very large libraries, as the number of constraints in the ILP formulation grows linearly with the number of selected mutants.

In future work, we plan to extend our method to consider the breadth optimization problem, where the goal is to design antibodies that are effective against a set of divergent viral strains. We also plan to investigate the use of a quadratic assignment formulation to model the pairwise interactions between amino acids in the antibody-antigen complex.

## Code availability

The code associated with this manuscript has been open-sourced to facilitate reproducibility and further research in the field. The repository contains the implementation of the *Solve-and-remove* Algorithm 1. It also includes scripts for data preprocessing and evaluation. The code is available at the following URL: https://github.com/LLNL/protlib-designer. Please refer to the repository’s README file for setup instructions, dependencies, and usage guidelines. For any questions or issues regarding the code, please contact the corresponding author.

## Acknowledgements

This work was performed under the auspices of the U.S. Department of Energy by Lawrence Livermore National Laboratory under contract DE-AC52-07NA27344. Lawrence Livermore National Security, LLC. LLNL-CONF-868784.

## A APPENDIX

### A.1 Results for Spesolimab

Below, we present the results of our experiments using the Spesolimab system. Table 3 contains the PLD results with all baselines evaluated using a number of metrics.

### A.2 Code Availability

The code accompanying this manuscript has been open-sourced to enhance reproducibility and support further research. For review, it is provided as a ZIP file in the supplementary material. Upon acceptance, a GitHub repository link will be included in the manuscript. The repository includes the implementation of Algorithm 1, along with scripts for data preprocessing and evaluation.

### A.3 Cold Start Antibody Library Design

We focus on a cold start setting for antibody library design, whereby no experimental or computational fitness data is required. While experimental wet lab data or computational physics-based simulators can be used to seed antibody library design, these approaches can be very expensive, time consuming, and require extensive computational resources. As a result, many of these tools are not accessible to the average researcher. Recent open sourced deep learning models (e.g. ESM, ProteinMPNN, AntiFold) have been shown to accurately predict the biological effects of mutations [51]. By leveraging these models as a surrogate fitness signal, our method offers a fast, low-cost route to generate diverse antibody libraries. Moreover, when experimental or physics-based fitness data are available, ProtLib-Designer can seamlessly incorporate them to further refine its designs.

### A.4 Search Space Definition for Trastuzumab, D44.1, and Spesolimab

For each system the size of the search space (all possible mutants) can be calculated as follows:

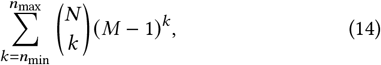

where *n*_min_ is the minimum number of mutations per sequence, *n*_max_ is the maximum number of mutations per sequence, *N* is the number of mutable positions, and *M* = 20 is the number of amino acids. For each system, *n*_min_ = 5 and *n*_max_ = 8.

Following the work of [34], we define the search space for the Trastuzumab antibody (PDB 1N8Z) as a set of 10 mutable positions on the CDR3 region of the heavy chain. The set of mutable positions contains the following positions: H99, H100, H101, H102, H103, H104, H105, H106, H107, H108. Using 14, the total number of possible mutants for Trastuzumab is ≈ 8 × 10^11^.

For the D441 antibody (PDB 1MLC), we follow [52] and consider a total of 34 mutable positions on the heavy and light chains. The set of mutable positions contains the following positions: H28, H30, H31, H37, H43, H47, H57, H59, H60, H69, H72, H73, H74, H76, H102, H104, H110, L22, L24, L25, L26, L28, L32, L37, L40, L43, L51, L52, L54, L55, L85, L89, L92, L93. With 34 mutable positions and using 14, the total number of mutants for D44.1 is ≈ 3 × 10^17^.

For the Spesolimab antibody (PDB 6U6U), we follow [46] and consider a total of 47 mutable positions spanning all heavy and light chain CDRs. The set of mutable positions is: H26, H27, H28, H29, H30, H31, H32, H33, H51, H52, H53, H54, H55, H56, H57, H58, H97, H98, H99, H100, H101, H102, H103, H104, H105, H106, H107, H108, L27, L28, L29, L30, L31, L32, L33, L51, L52, L53, L90, L91, L92, L93, L94, L95, L96, L97, L98. Using 14, the total number of mutants for Spesolimab is ≈ 5 × 10^18^.

### A.5 Antibody Library Design as a Multi-Objective Problem

Designing a library with respect to *extrinsic fitness*, e.g. binding quality to the antigen target, does not ensure experimental success. Safe, stable, and manufacturable antibody candidates contain key properties related to *intrinsic fitness* e.g thermostability, developability, and stability. Without explicitly considering the *intrinsic fitness* related properties during optimization a generated library may contain candidates that overfit to the biases of the optimized *in silico* tool increasing the risk of experimental failure. As a risk mitigation strategy, we propose to optimize for *intrinsic fitness* and *extrinsic fitness* simultaneously as separate objectives.

Approaches that frame antibody library design as a multi-objective problem typically compute a Pareto front of solutions, or optimize the library for a fixed weighting over the problem objectives [9]. As we will show, the Pareto front does not contain sufficiently diverse solutions from which an adequate antibody library can be derived. Moreover, in a *zero-shot* setting, optimizing a library with respect to a fixed weighting over the objectives increases the risk of experimental failure. Objective weightings are extremely difficult to tune [17] and, as a further complicating factor, it is infeasible to determine if any selected weighting is appropriate given the absence of experimental fitness data. We utilize a dynamic weighting approach [1], whereby for each iteration a random weighting over the objectives is sampled from the distribution over all possible weightings, and used to compute a feasible solution for the given problem. Sampling weights mitigates the risk of over optimizing for any individual weighting, ensuring diversity and coverage over the space of objective weights.

### A.6 Tunable hyperparameters for ProtLib-Designer

In table 4 we present the tunable hyperparameters for ProtLib-Designer. The small number of tunable parameters presented in table 4 ensures researchers can carefully design their experiments with respect to specific design considerations without extensive hyperparameter tuning.

**Table 4:** List of tunable parameters for ProtLib-Designer.

| Parameter Name | Symbol |
| --- | --- |
| Positional diversity constraint | $\delta_1$ |
| Mutational diversity constraint | $\delta_2$ |
| Ball radius | $\epsilon$ |
| Cyclic mutation count | $\xi$ |

### A.7 Baseline Algorithms

Below we outline the LMG, SPEA2 and MODIFY algorithms, used as baselines to compare against ProtLib-Designer. We also describe how an approximation of the Pareto front is computed, which we also use as a comparison.

#### Linear mutant generator

The Linear mutant generator (LMG) baseline is adapted from [8]. The LMG uses a hierarchical sampling process. First, the number of mutations is sampled from the range [5, 8] uniformly at random. Then, mutations are sampled without replacement according to the following probabilities,

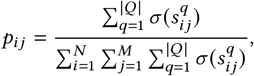

where σ (*s*) is the generalized logistic function 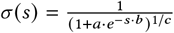, with *a* = 1000, *b* = 5, and *c* = 8. The LMG algorithm can be used to generate a batch of *K* mutants.

#### Strength Pareto Evolutionary Algorithm 2

As a baseline, we implemented the Strength Pareto Evolutionary Algorithm 2 (SPEA2) [58], enhanced with an island model strategy. SPEA2 was chosen for its balance between convergence and diversity, crucial in exploring the complex design space of antibody optimization. For the Trastuzumab experiment, we used an island model with 20 independent sub-populations, each with 50 individuals, resulting in a total of 1,000 individuals across the entire population. Each sub-population evolved independently, with no migration, ensuring isolated genetic pools and promoting diverse exploration of the solution space. The number of generations was set to 500. For the D44.1 system, we employed 10 sub-populations, each with 100 individuals, and ran SPEA2 for 1,000 generations.

A restart strategy was introduced to maintain diversity. If a sub-population’s Pareto Front (PF) was a subset of another subpopulation’s PF, the sub-population was restarted by replacing all individuals with random solutions. This mechanism helped prevent premature convergence within any sub-population, encouraging ongoing exploration. The restart mechanism was disabled for the last 10% of generations to prevent random solutions from appearing in the final population.

In our SPEA2 implementation, each solution (antibody mutant) is represented as y = (*y*_1_, …, *y*_*N*_ ), where *y*_*i*_ = ℐ_*A*_ (*x*_*i*_) and *x*_*i*_ ∈ *A*. Here, *y*_*i*_ indicates the index of the amino acid selected for the *i*-th mutable position from the set of amino acids *A*, and *N* represents the number of antibody positions permitted to mutate.

The crossover operator was a modified version of one-point crossover that respected the constraint of 5 to 8 amino acid mutations per antibody sequence. After the traditional one-point crossover was applied, if the offspring exceeded the upper mutation limit, random mutations were reverted to the wild-type sequence. If the number of mutations fell below the lower threshold, additional random amino acid mutations were introduced. The probability of crossover was set to 70%.

A custom mutation operator was also used, which randomly mutated an amino acid at a position that already had a mutation to a different amino acid. This ensures the total number of mutations remained constant and within the specified range. The probability of mutation was 30% at the individual level and 5% at the position level. The algorithm ran for 500 generations, with the final population comprising all unique individuals of the joint sub-populations, ensuring a broad and diverse set of solutions. We used the SPEA2 implementation from the DEAP [14] Python package.

In our experiments, we used two SPEA2 setups called *stratified* and *unified*. In the *stratified* setup, we ran four separate SPEA2 instances, each constrained to find solutions with an exact number of mutations (5, 6, 7, or 8). The final antibody library was composed of all unique solutions from the combined selections of these instances, ensuring diversity in the number of mutations. In the *unified* setup, a single SPEA2 instance was run across the entire mutation range [5, 8].

#### ML-optimized library design with improved fitness and diversity

The *ML-optimized library design with improved fitness and diversity* algorithm (MODIFY) was introduced in [9], following the work in [56]. The method optimizes a tensor ϕ ∈ ℝ^*N* ×*M*^ and uses it to build probabilities 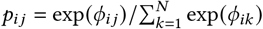 for each mutable position *i*. The probability of a mutant is given by

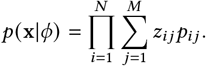

The optimization objective is,

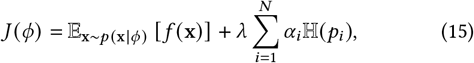

where 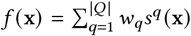 is the weighted sum of the scores of the |*Q*| objectives, *w*_*q*_ is the weight of the *q*-th objective, *s*^*q*^ (x) is the score of the *q*-th objective, and ℍ (*p*_*i*_) is the entropy of the marginal distribution *p*_*i*_ over amino acids at position *i*. The method sweeps through 200 λ values, optimizes the tensor ϕ for each λ value using policy gradient, and then selects the best λ value based on the area under the expected fitness and entropy curve. Objective (15) has been extensively used in other works for discrete optimization [31, 39].

Here, we adopt the MODIFY algorithm with *Q* = {PLM, IFOLD}, the scores defined in (3)-(8), and, as suggested in [9], *w*_PLM_ = *w*_IFOLD_ = 0.5. We train the models for 2, 000 iterations, with a batch size of 1, 000, and a learning rate of 10^−1^. The final library of 1, 000 mutants was obtained by sampling from the optimal distribution *p* (x|ϕ_λ=0.66_) (see Figure 5).

**Figure 5:**
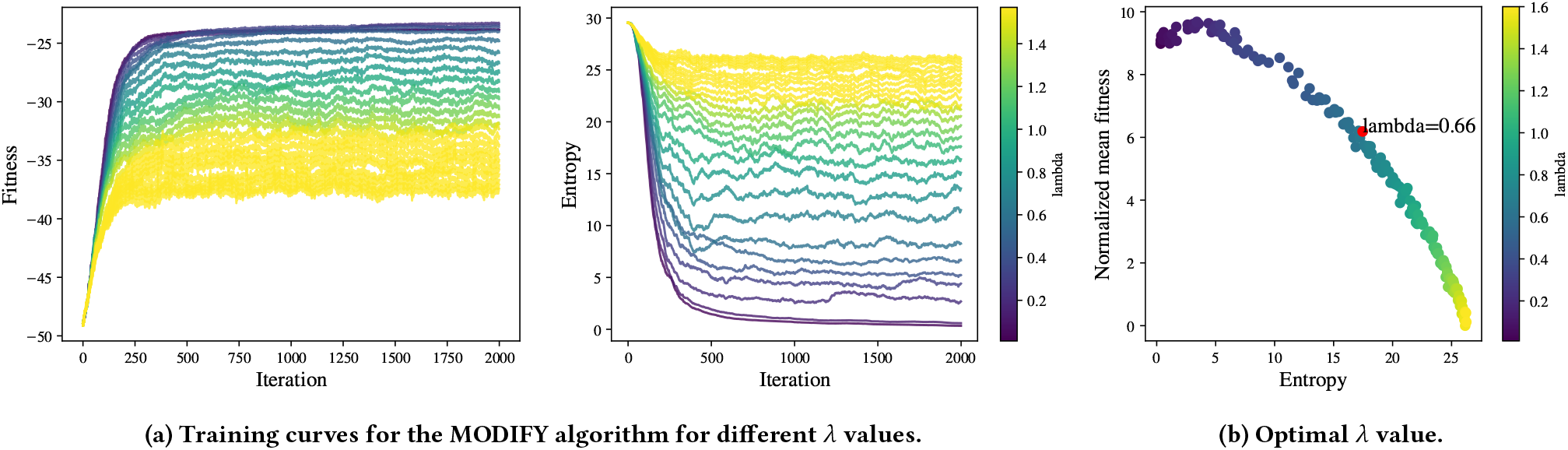
(a) Evolution during training of the fitness and entropy of the MODIFY algorithm for Trastuzumab. (b) The optimal λ value maximizes the area under the curve of the expected fitness and entropy.

### A.8 Approximating the Pareto Front

It is infeasible to exhaustively compute all valid solutions over the search space for 5 − 8 point mutations. As a result, exactly computing the Pareto front is difficult without access to an exhaustive set of solutions. Therefore, we compute an approximation of the Pareto front by first computing the Convex hull of the Pareto front, then running Pareto optimality filter on the combined sets of the Convex hull with ProtLib-Designer, SPEA2, MODIFY, and LMG runs. The resulting set of solutions is then used as the approximation of the Pareto front shown in the main text.

The Convex hull is a subset of the Pareto front [17, 41]. ProtLib-Designer utilizes an optimal solver, meaning for any linear weighting ProtLib-Designer can find the optimal solution that lies on the Convex hull. To compute the Convex hull, we run ProtLib-Designer without the solve-and-remove algorithm, therefore the solution landscape remains static during optimization. To uncover the Convex hull, at each iteration we sample a different linear weighting and solve with respect to the sampled weighting. For our experiments, we sampled a total of 1,000 weightings. The resulting Convex hull is presented in fig. 6. While the Convex hull contains many solutions that lie on the Pareto front, it does not contain solutions that lie on the concave regions of the Pareto front. Therefore, we initiate another step to approximate the remaining solutions in these missing regions.

**Figure 6:**
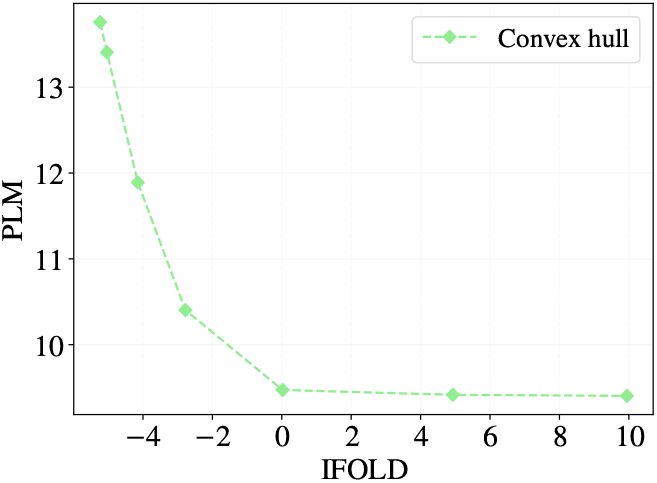
Convex hull computed using ProtLib-Designer without the solve-and-remove algorithm for Trastuzumab. Generated using 1, 000 sampled linear weights.

To find the missing regions, we first compute the Pareto fronts of the solution set computed by each algorithm. In fig. 4a, it is clear that the solutions computed by the LMG and MODIFY algorithms are all Pareto dominated by the SPEA2 and ProtLib-Designer Pareto fronts. Both ProtLib-Designer and SPEA2 runs compute identical Pareto fronts, which also contain all solutions on the Convex hull, and solutions in concave regions. As a result, we use the resulting Pareto front as our approximation presented in the experimental section.

### A.9 Evaluation Metrics

To measure each individual batch produced by the ILP, MODIFY, LMG, and SPEA2 algorithm we utilized a number of metrics to compare the performance of each algorithm.

#### Evaluating extrinsic fitness

To measure the extrinsic fitness of a batch of mutated sequences, we employ an oracle function to approximate the binding potential of a given mutated sequence to the target antigen. The oracle is a trained predictive classification model and will act as a surrogate for experimental validation. To train the oracle, we utilized extensive experimental binding data from [34]^2^. The corresponding dataset contains 31, 000 experimentally validated and labelled mutated sequences, where 11, 000 bind to the target antigen (labelled as 1) and 20, 000 do not bind to the target antigen (labelled as 0). During training we use a random 70 /10 /20 train/val/test split with binary cross-entropy loss for 20 epochs and a batch size of 16. The trained oracle network achieves 83% accuracy on the test set. To measure the extrinsic fitness of each batch, we report the percentage of sequences in the batch that are predicted by the oracle to bind to the target.

#### Evaluating intrinsic fitness

Similarly to Gruver et al. [16], to measure the intrinsic fitness of a batch of mutated sequences we utilize the log-likelihood assigned by ProtGPT2 [11] (trained on Uniref50 [50]). The resulting score for each sequence represents how likely a given sequence is with respect to the model. The loglikelihood assigned by ProtGPT2 can be interpreted as a humanness score, where a lower score corresponds to a sequence being more “natural” or human-like. Therefore, we refer to this measure as humanness in the experimental section. To calculate the humanness of a given batch, we compute the mean over the batch with respect to the humanness score.

#### Diversity metrics

We use the residue entropy to measure the diversity of the batch. The residue entropy refers to the entropy of the empirical distribution over amino acids and positions of a batch, *B*. Consider the empirical distribution, 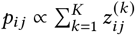, the residue entropy of the batch is defined as

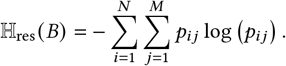

As entropy increases a distribution becomes uniform. As a result, we aim to maximize entropy to compute a batch that is uniform over possible amino acids and mutable positions, therefore making the batch more diverse.

#### Multi-objective metrics

To measure the ability of each algorithm to optimize each objective we utilize the *hypervolume* metric and a *based expected utility* metric.

The hypervolume (HV) metric measures the volume in vector space of a given Pareto front, which correlates to the spread of a given undominated set over the possible multi-objective solutions space. The HV is defined as

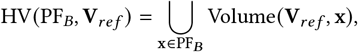

where Volume (**V**_*ref*_, x )is the volume of the hypercube spanned by the reference vector, **V**_*ref*_, and the solution vector x in the Pareto front of a given batch, PF_*B*_ [17]. To measure the HV in our experiments we set **V**_*ref*_ = [50, 50] .

While the hypervolume metric has been used extensively in the literature, it has some limitations. Specifically, the hypervolume metric requires a reference point to compute the score. The choice of reference point is arbitrary and affects the resulting score.

Many multiobjective measures focus on evaluating an algorithm’s ability to compute a coverage set of non-dominated solutions, e.g. Pareto front [41]. However, for antibody library design, we are concerned with measuring the quality with respect to the objective values over a batch of solutions. To do so, we implement a *batch expected utility* metric (BEU) that takes advantage of the expected utility metric (EUM) proposed by Zintgraf et al. [57]. The BEU metric produces a scalar representation of the expected utility over a given batch of solutions with respect to a distribution over utility functions, and can be defined as follows:

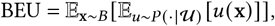

where *u* is a utility function drawn from a set of utility functions 𝒰, and x is a mutant in the batch, *B*.

To compute the BEU metric, a distribution over utility functions must be known a priori. Therefore, we assume a distribution over linear utility functions and sample from a simplex over all possible convex combinations. In our experiments, we sample 10, 000 utility functions (linear scalarization weights) and compute the BEU metric for each algorithm in tables 1 to 3.

### A.10 Evaluating Deep Learning Scoring Functions

#### A.10.1 Correlations Between Experimental Data and In-Silico Tools

The available data in [52] for D44.1 contains experimental wet lab data for single point mutations in the wild-type antibody. The experimental single point mutations are scored using a negative log enrichment. This scoring mechanism is commonly used in protein design and quantifies how enriched or depleted a mutant is. Furthermore, the authors in [22] calculated the FoldX ΔΔ*G* scores for all single point mutations and made that data available. FoldX [43] is a computational simulation tool that estimates the change in free energy (ΔΔ*G*) caused by point mutations in a protein structure.

To understand how experimental wet lab data and computational simulation tools correlate with state-of-the-art deep learning tools, we generate scores for a number of deep learning tools and compare the correlation between each tool, the FoldX ΔΔ*G* values, and the experimental single point negative log enrichment data. The deep learning tools that we compare are ESM, ProtBERT, and AntiFold.

Figure 7 displays the correlation matrix of the various scores. The deep learning models—ESM, ProtBERT, and AntiFold—show moderate positive correlations (0.35–0.46) with the negative log enrichment values, indicating that, despite being trained purely by maximum likelihood, these methods capture meaningful aspects of a mutation’s biological impact. In contrast, FoldX scores correlate negatively with the same enrichment metric. Notably, ESM and ProtBERT also exhibit a modest positive correlation with FoldX. Overall, each of the deep learning tools outperforms FoldX in terms of correlation strength, and the three deep learning models themselves are strongly intercorrelated.

**Figure 7:**
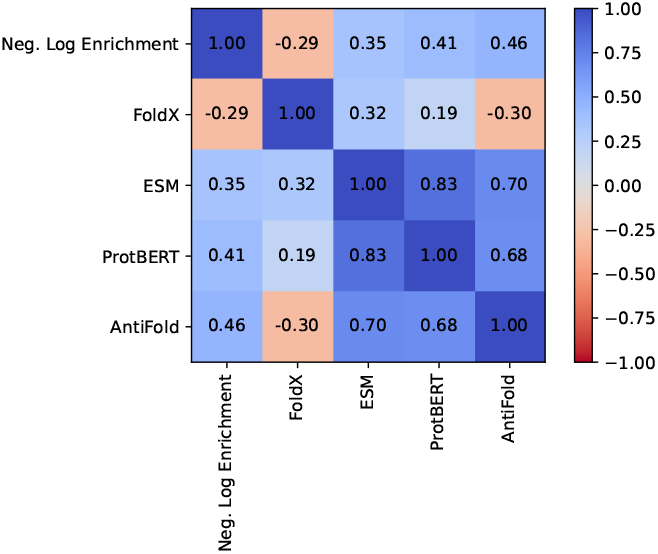
Correlation matrix for D44.1 scoring methods for wet lab, computational and deep learning tools that quantify mutational effects.

It is important to emphasize that these findings pertain solely to the D44.1 system; correlations may differ in other contexts. A more exhaustive evaluation would entail applying this analysis across multiple antibody–antigen complexes—ideally spanning various viral families—and comparing additional simulation platforms (for example, Rosetta) alongside other machine-learning approaches (such as ProteinMPNN). An example of such evaluation can be found in [51].

#### A.10.2 Deep Learning Tools and AlphaFold3

To further investigate the use of deep learning tools as optimization signals during our experiments, we generate a library of 1, 000 sequences for D44.1 using the ProtLib-Designer while optimizing ProtBERT and AntiFold scores and evaluate each generated library under AlphaFold3 (AF3) [2].

AF3 reliably predicts antibody–antigen complex structures and provides per-prediction confidence metrics. In particular, recent work [20] has demonstrated that a high heavy-chain interchain predicted TM-score (ipTM *>* 0.8) correlates well with DockQ, a standardized measure of docking accuracy earned by comparing predicted and experimental complexes. Thus, heavy-chain ipTM is an interesting metric to compute, as it can be used as a proxy for the structural success of a given mutant.

We executed ProtLib-Designer in two modes—one optimizing AntiFold, the other ProtBERT—each using the solve-and-remove algorithm. Figures 8 and 9 summarize the outcomes. For each variant in the 1,000-sequence library, AF3 was run with 50 random seeds, and we report the highest heavy-chain ipTM across those seeds. To show how ProtLib-Designer steered sequence selection, we also plot each variant’s negative log-likelihood ratio (NLL) from the deep learning model(s). Point color indicates the mutation rank (i.e., the order in which ProtLib-Designer selected that mutant), while point shape encodes its Hamming distance from the D44.1 wild type.

**Figure 8:**
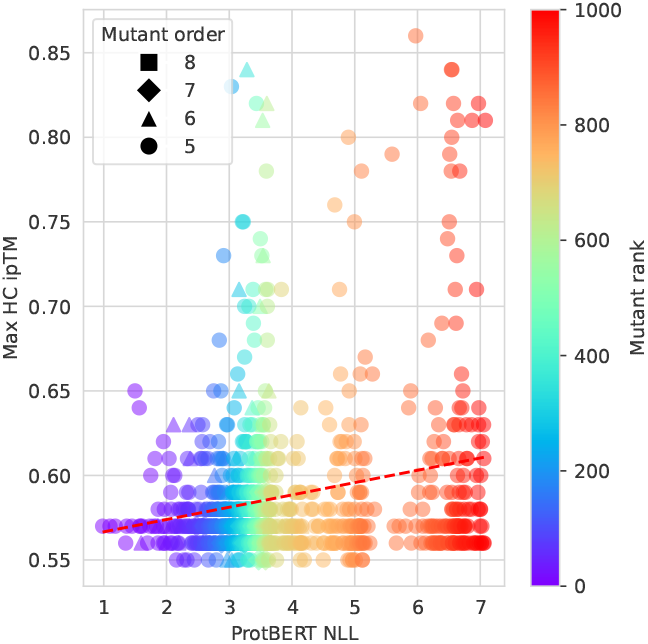
Correlation between ProtBERT NLL and AF3 HC ipTM for the mutated sequences generated by ProtLibDesigner. The color of each point represents the order in which the mutant was selected by ProtLib-Designer (purple = first selected, red = last selected). The shape of each point represents the number of mutations from the D44.1 wildtype.

**Figure 9:**
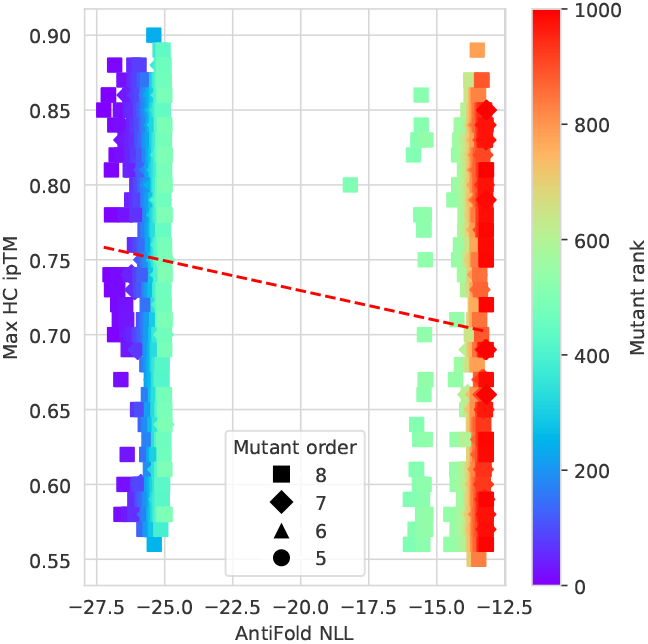
Correlation between AntiFold NLL and AF3 HC ipTM for the mutated sequences generated by ProtLib-Designer. The color of each point represents the order in which the mutant was selected by ProtLib-Designer (purple = first selected, red = last selected). The shape of each point represents the number of mutations from the D44.1 wildtype.

#### ProtBERT

Figure 8 plots each ProtLib-Designer variant’s ProtBERT NLL against its maximum heavy-chain ipTM from 50 AF3 runs. While ProtLib-Designer explores lower-NLL sequences first, there is little relationship between ProtBERT score and ipTM—and only a handful of mutants exceed the 0.8 ipTM threshold. In this system, ProtBERT alone appears to be a weak guide for structural quality.

#### AntiFold

By contrast, optimizing AntiFold NLL yields mutants that span the full range of ipTM values, with many surpassing 0.8 (Figure 9). Because AntiFold conditions on both antibody and antigen, its NLL more reliably flags mutations that improve or at least maintain predicted interchain packing.

#### Conclusions

AF3’s heavy-chain ipTM can serve as a useful proxy for structural consistency in antibody–antigen complexes, but it comes at substantial computational cost (typically requiring a GPU cluster, computing multiple sequence alignments, and running multiple seeds). In our comparison, ProtBERT scores showed little predictive power, while AntiFold’s context-aware NLL more reliably flagged variants with higher ipTM. By using deep-learning signals—especially those that incorporate antigen context—to triage candidates before expensive AF3 runs, we can both cut computational expense and enrich for structurally promising mutants.

### A.11 ProtLib-Designer Parameter Ablations

Below we outline a number of ablations using ProtLib-Designer with Trastuzumab.

In table 5, we show results of ablations of ProtLib-Designer experiments for Trastuzumab with different diversity constraints and cyclic mutation count parameter configurations. In all experiments, the Ball radius parameter ε was set to zero to enforce that no duplicate mutants are selected, ensuring basic non-redundancy in the designed library and allowing better investigation of the effect of the other parameters. In table 5, fitness is measured as the percentage of the generated library predicted to bind the antigen HER2 target using the binding prediction surrogate. For experiment 1, we evaluate ProtLib-Designer without any diversity constraints and without cyclic mutation count parameter (ξ = 0) enabled. In this case, ProtLib-Designer is greedy and selects sequences only with respect to the weighted AntiFold and ProtBERT scores, and achieves a fitness of 61.8%. For evaluations 2-6, we introduce diversity constraints, and vary them for each experiment respectively. We focus on varying the mutational constraint, δ_2_, from 500 to 100, to evaluate the effect of this constraint on the fitness and diversity of a given batch. For δ_2_ = 500 and δ_2_ = 400 a small change in the fitness and diversity of the batch is observed. However, as the δ_2_ constraint is reduced further (δ_2_ *<* 400) the predicted fitness of each batch decreases. Simultaneously, as the δ_2_ constraint is reduced, the entropy of each batch increases resulting in more diverse batches. A trade-off between predicted fitness and diversity is observed, where generating batches of higher diversity results in a lower predicted fitness.

**Table 5:** Ablation of ProtLib-Designer experiments with various diversity parameters and the resulting predicted fitness and entropy scores for Trastuzumab.

| Exp. id | Diversity constraints | $\delta_1$ | $\delta_2$ | $\xi$ | Entropy | Pred. Fitness (%) |
| --- | --- | --- | --- | --- | --- | --- |
| 1 | × | NA | NA | 0 | 3.11 | 61.8 |
| 2 | ✓ | 1000 | 500 | 0 | 3.10 | 62.9 |
| 3 | ✓ | 1000 | 400 | 0 | 3.12 | 66.2 |
| 4 | ✓ | 1000 | 300 | 0 | 3.22 | 58.2 |
| 5 | ✓ | 1000 | 200 | 0 | 3.47 | 44.5 |
| 6 | ✓ | 1000 | 100 | 0 | 4.02 | 28.5 |
| 7 | × | NA | NA | 1 | 2.96 | 40 |
| 8 | ✓ | 1000 | 500 | 1 | 3.09 | 39.7 |
| 9 | ✓ | 1000 | 400 | 1 | 3.21 | 37.4 |
| 10 | ✓ | 1000 | 300 | 1 | 3.33 | 32.8 |
| 11 | ✓ | 1000 | 200 | 1 | 3.66 | 22.9 |
| 12 | ✓ | 1000 | 100 | 1 | 4.26 | 13.4 |

fig. 11a visualizes the scores for the problem objectives for each mutated sequence computed by ProtLib-Designer experiments, where experiments 1 and 4 are shown in fig. 11a from table 5. Additionally, we compute the Pareto front, represented in fig. 11 as a dashed line. Both ProtLib-Designer configurations find all sequence mutations on the Pareto front. Experiment 1 (red), the ProtLib-Designer instance without diversity constraints, greedily selects sequences close to, or on, the PF. It is clear, the solve-and-remove algorithm (algorithm 1) enables some level of diversity even without diversity constraints being explicitly configured. Without the solve- and-remove algorithm, only solutions on the PF would be selected. In contrast, experiment 4 (teal), the ProtLib-Designer instance with diversity constraints, produces a batch of sequences where some solutions lie close to the PF, but many solutions span different regions of the objective space, resulting in more pronounced diversity. The diversity for experiment 4 is visualized as a sequence logo in fig. 10b. The entropy of experiment 4 is 3.22, and is higher (i.e., more diverse) when compared to the entropy score of 3.11 for experiment 1.

**Figure 10:**
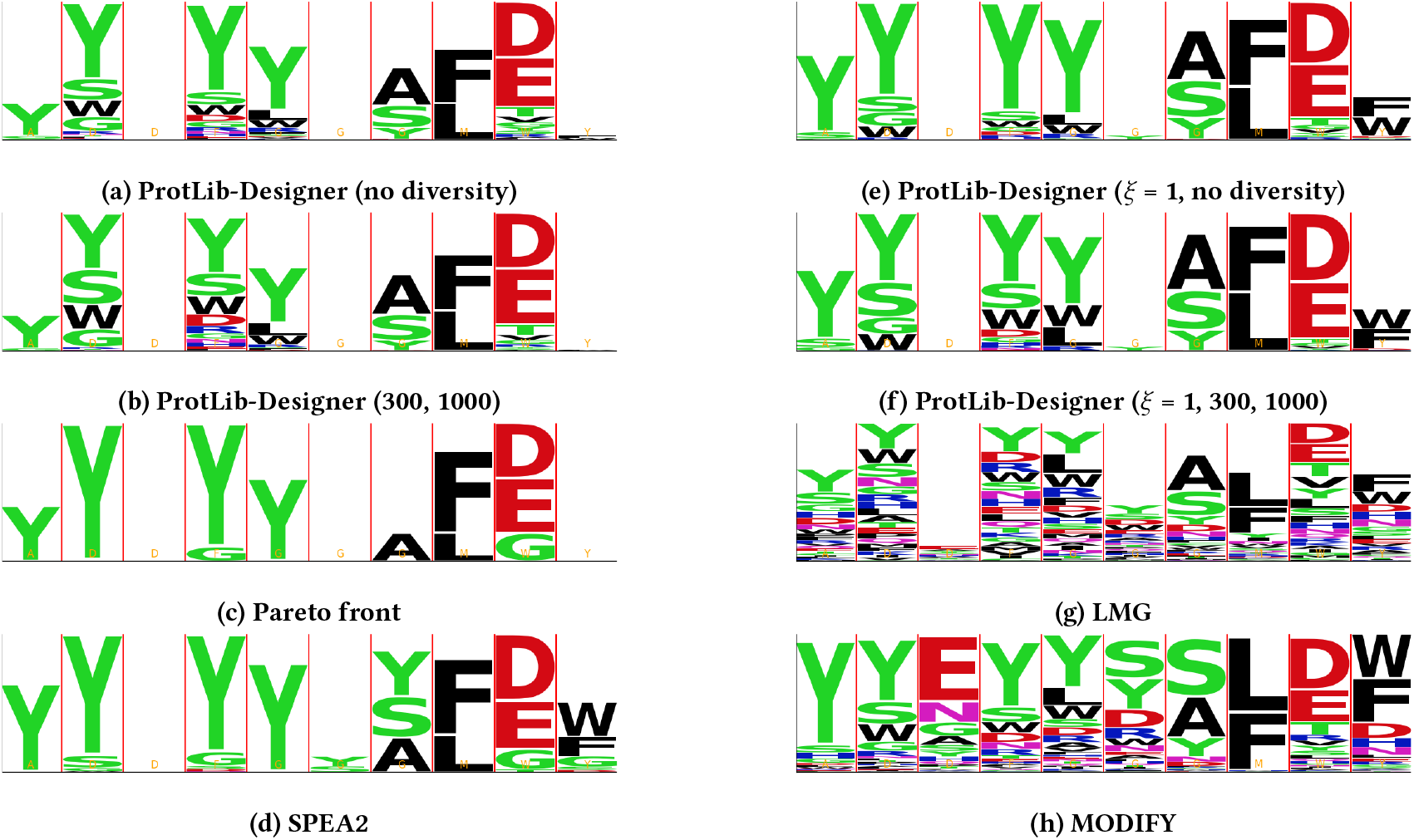
Sequence logo plots of batches generated from various experiments for ProtLib-Designer applied to the Trastuzumab system, each respective baseline, and the Pareto front.

**Figure 11:**
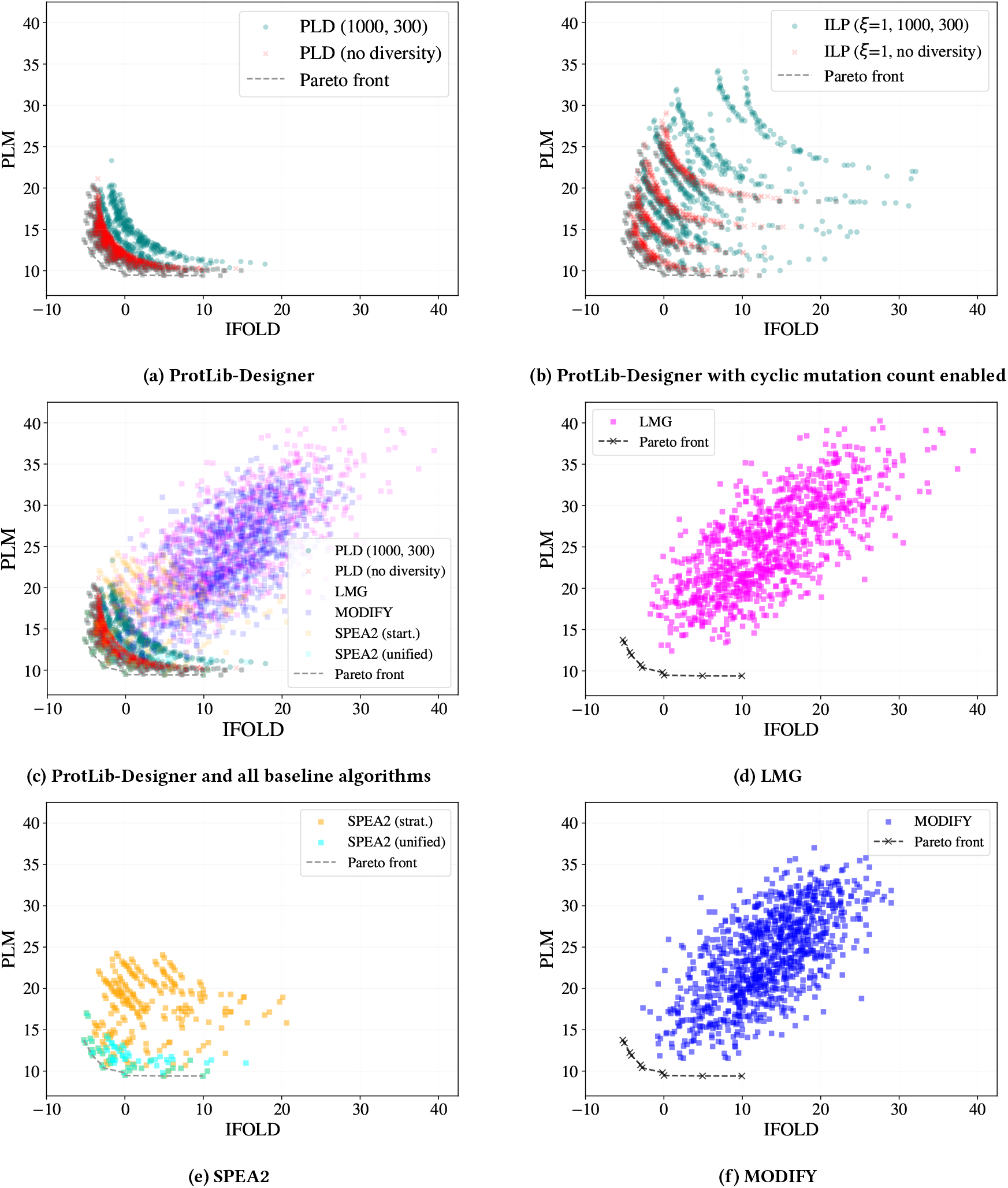
The objective values of the batch of sequences generated by ProtLib-Designer, LMG, SPEA2, and MODIFY algorithms. Each point corresponds to a unique mutated sequence.

Next, we investigate the impact of the cyclic mutation count parameter (ξ = 1) on the batches generated by ProtLib-Designer. For experiments 7-12, we introduce the cyclic mutation count parameter with and without diversity constraints. Notably, the diversity of each batch with the cyclic mutation count parameter enabled increases when compared to the corresponding experiments without the cyclic mutation count parameter. fig. 11b presents the objective scores for experiment 7 and 10, where the objective score values are distributed differently when compared to their corresponding experiments in fig. 11a. For example, the solutions generated in experiment 7 are much further from the PF when compared to the solutions generated in experiment 1 which lie close to the PF. Additionally, the solutions generated in experiment 10 are scattered throughout the objective space. As a result, the cyclic mutation count parameter impacts the fitness of each batch, given all experiments with the cyclic mutation count parameter have a lower fitness compared to their corresponding experiment. The cyclic mutation count parameter ensures more diversity in the point mutations of the given batch and without the cyclic mutation count parameter enabled this level of diversity cannot be guaranteed.

### A.12 Further Baseline Comparisons and Discussion

Below we extend our comparison of ProtLib-Designer runs against with LMG, MODIFY, and SPEA2. We also use an approximation of the Pareto front for further comparisons. We focus on Trastuzumab, however, similar conclusions can be drawn for D44.1.

#### Unique sequences

Library design typically requires a fixed number of sequences to be generated to fit a capacity for experimental validation. Therefore, it is important to have control on the number of unique sequence generated by a library design algorithm. Using ProtLib-Designer, MODIFY, and the LMG it is possible to generate a library of fixed size, given each algorithm generates the required library size of 1, 000. However, the SPEA2 algorithm struggles to generate batches that meet this requirements. SPEA2 (joint) generates a batch of 69 unique sequences, while SPEA2 (stratified) generates a batch of 263 sequences. Although the initial population size of the SPEA2 runs was set to 1, 000 many similar, or the same solutions, were generated or dropped due to the algorithms optimization process, resulting in smaller batch sizes. As a result, the inability to control the exact size of the library being generated limits the SPEA2 algorithms use in practical settings.

#### Intersection of solution sets

Figure 12 shows UpSet plots [32] illustrating the intersections of solution sets obtained by ProtLib-Designer and baseline methods for trastuzumab and D44.1. In general, we observe limited overlap between the solution sets. Specifically, there is minimal to no intersection between LMG and MODIFY with other methods, while there is a small overlap between the top-performing methods, SPEA2 and ProtLib-Designer. For trastuzumab, 50 variants are shared between ProtLib-Designer and SPEA2-Stratified, and for D44.1, 250 variants are shared. Although the full solution sets differ, the Pareto sets—comprising solutions on the Pareto Front—are identical for the top performers, ProtLib-Designer and SPEA2, as noted earlier. This suggests that SPEA2, while effectively searching for the optimal Pareto set, lacks a mechanism to explore solutions near the Pareto set, unlike ProtLib-Designer, which targets both the Pareto Front and nearby solutions.

**Figure 12:**
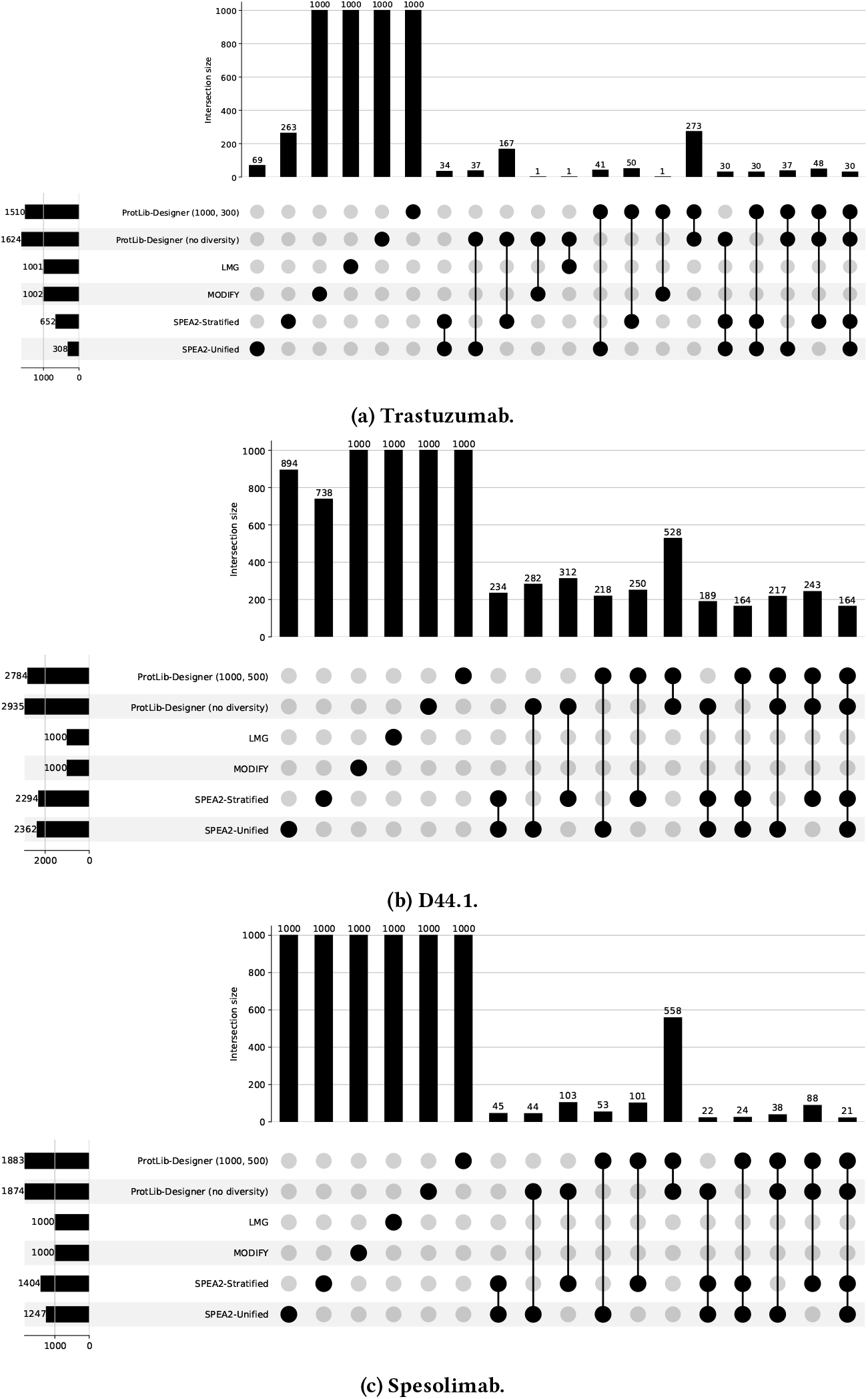
Intersection of solution sets across all the methods investigated.

#### Residue entropy and oracle fitness

The measured entropy of the distribution over amino acids and positions is utilized score the diversity of a given batch, where a diverse batch of sequences maximizes the entropy of the distribution. We also use the predicted oracle fitness of the batch using a trained predictive model (see the experimental section) to measure the extrinsic fitness of each batch. The measured entropy of the LMG generated library is 4.75, resulting in a highly diverse batch of solutions. Similarly, the MODIFY algorithm has a residue entropy score of 4. Both LMG and MODIFY achieve higher diversity scores when compared to ProtLib-Designer experiments presented in table 5. fig. 10g and fig. 10h display the sequence logo plot of the batch produced by LMG and MODIFY. The difference in diversity obvious when comparing the MODIFY and LMG logo plots with those of ProtLib-Designer runs (fig. 10a, fig. 10b, fig. 10e, and fig. 10f). Each position contains a number of amino acids, resulting in a highly diverse batches. However, LMG and MODIFY have lower predicted oracle fitness scores, 17.1 and 18.6, when compared to ProtLib-Designer runs (61.8 and 58.2) in table 1.

Both SPEA2 runs also have lower diversity scores when compared to ProtLib-Designer runs. SPEA2 (unified) has a score of 2.6 and SPEA2 (stratified) has a score of 2.74. This is reflected in fig. 10d, where many amino acids are missing from the logo plot at each mutated position. The SPEA2 (unified) has a fitness score of 68.12 and SPEA2 (stratified) has a fitness score of 68.12. Both SPEA2 runs have a lower diversity score when compared to ProtLib-Designer runs. However, the SPEA2 (stratified) run has a higher fitness score than ProtLib-Designer. The SPEA2 (stratified) batch only has 69 sequences, where 10 of those sequences lie on the Pareto front. Therefore, the results from the SPEA2 runs are not directly comparable. ProtLib-Designer produces a much larger batch when compared to both SPEA2 runs and maintains a high level of both diversity and fitness.

Given that multiple objectives are used during optimization, we approximate the Pareto front of the underlying problem and compare the set of sequences in the Pareto front with ProtLib-Designer and LMG. The set of sequences on the Pareto front have a high fitness (80%) but a very low diversity (2.21). Although the Pareto front has fit solutions, the Pareto front is not sufficiently diverse to use as a batch for experimental validation. However, we should aim to include the Pareto front as a subset of solutions contained with the overall batch.

The diversity parameters for LMG and MODIFY are not configurable, resulting in algorithms that always produce batches with high diversity. Similarly for SPEA2, certain parameters can be altered to enforce diversity; however, the diversity of the resulting set is not directly controllable. In contrast to all other baselines, ProtLib-Designer can be configured to generate more or less diversity within a batch when required. Given that various library design scenarios can arise where varying levels of diversity may be necessary, having the ability to configure the diversity parameters as required is important.

fig. 13 presents the trade-off between fitness and diversity with respect to ProtLib-Designer and each baseline. Each baseline is shown as a single point to reflect the fitness and diversity score of the final batch. ProtLib-Designer contains multiple points representing each ablation performed. It is clear, that to increase fitness scores the diversity of the batch must decrease. As diversity increases, the fitness of the batch also declines. Therefore it is important to have control over the diversity of a batch, given depending on the objectives certain diversity parameters may be required.

**Figure 13:**
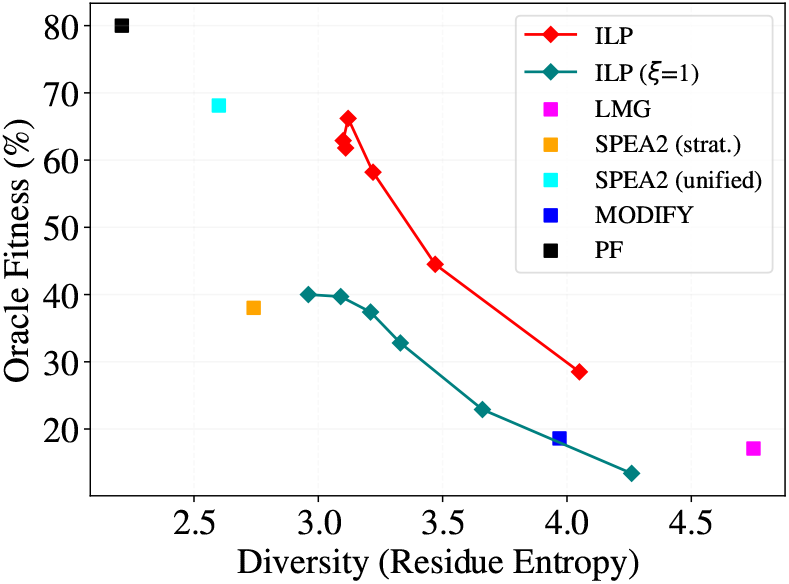
Predicted oracle fitness and residue entropy based diversity of each ProtLib-Designer (PLD) ablations against the baseline algorithms.

#### Humanness

While the typical objective of library design is to generate antibodies that bind to the given target, it is also crucial to ensure that a given antibody library contains sequences with properties related to *intrinsic fitness* such as developability, humanlikeness, etc. – to ensure experimental success. Sequences without properties related to *intrinsic fitness* may not be developable, human-like, or stable. To generate a library with these properties, we optimize the PLM scores as previously outlined. We measure the humanness of a given batch using the log-likelihood scores from a PLM, specifically ProtGPT2. The LMG and MODIFY generate libraries with scores of −1309.75 and −1308.27. In comparison, ProtLib-Designer runs have humanness scores of −1293.68 and −1296.73. Furthermore, the SPEA2 algorithm produces batches with −1290.01 and −1282.57. We aim to maximize the humanness score of a given batch. The LMG and MODIFY scores perform worse compared to the ProtLib-Designer and SPEA2 scores. Again, it is important to note that the SPEA2 batches contain fewer sequences when compared to all other baselines, and average humanness scores for a given batch are not directly comparable. However, given the scores of both ProtLib-Designer and SPEA2 batches, both sequence libraries are more likely to contain *intrinsic fitness* properties related to developability compared to LMG and MODIFY. A library designed to contain developable (human-like) sequences may avoid biases or failure modes that can arise when optimizing specifically for binding. A resulting library may be more likely to avoid experimental failure by ensuring that key properties related to intrinsic fitness are contained within the library.

#### Hypervolume

To understand how effectively each objective is being optimized, we compute and compare the hypervolume (HV) of the Pareto front of the batch of solutions computed by the ILP and all baselines. The Pareto fronts computed by all algorithms are presented in fig. 4a, where the LMG (HV = 1937.7) and MODIFY (HV = 2012.43.7) algorithms struggle to compute solutions that effectively minimize the objectives. Both ProtLib-Designer and SPEA2 runs uncover the full approximated Pareto front which contains the Convex hull (see fig. 6), resulting in an equal HV score of 2231.90. ProtLib-Designer and SPEA2, contain good approximations of the Pareto front, given they both contain the computed Convex hull. Therefore, both ProtLib-Designer and SPEA2 effectively optimize the problem objectives.

#### Expected utility of the batch

To further measure the quality of each batch with respect to the defined objectives we compute the *batch expected utility* (BEU) score for each algorithm. Here, we aim to minimize the BEU because a lower BEU score corresponds to a batch that effectively minimizes the objective values across a range of utility functions. Both LMG (BEU = 10.7) and MODIFY (BEU = 10.14) achieve a similar BEU score. However, the SPEA2 and ProtLib-Designer BEU scores are much lower when compared to the LMG and MODIFY. The batch produced by the SPEA2 (unified) algorithm achieves the lowest BEU score, however, as previously stated, the batch contains only 69 sequences making the BEU score for SPEA2 runs not directly comparable. ProtLib-Designer runs achieve slightly higher scores of 4.30 and 4.612, however these scores are achieved over much larger batches. As expected, the Pareto front has the lowest BEU score.

#### Diversity of the Pareto front

Many multi-objective optimization methods aim to compute the Pareto front [17, 41], where the Pareto front is the set of Pareto non-dominated solutions. Pareto dominance enforces diversity in the objective space, given in the Pareto front no two solutions can be the same, and, by the nature Pareto dominance, the objective values within the set span the range of feasible solutions. Figure 10c displays the positional and mutational diversity of the Pareto front some positions contain no mutations while others contain little mutational variation. Additionally, the PF has a lowest measured entropy value of 2.21. Given the lack of diversity of the set of solutions on the Pareto front it is important to explicitly optimize for diversity in sequence space. As a result, solutions that may be Pareto dominated can potentially be included in the final batch given these solutions may promote diversity.

#### Computational cost of ProtLib-Designer

Figures 14 illustrate the computational cost of running ProtLib-Designer to generate a library of 1,000 sequences with constraints and the solve-and-remove algorithm enabled. In this setup, constraints are incrementally added at each iteration, and the solver must account for all prior constraints when generating a new solution. This results in a non-linear increase in CPU time as the number of iterations grows. Without constraints or the solve-and-remove strategy, the computational cost would remain relatively constant across iterations. However, as shown in both figures, the CPU time per iteration increases significantly as more constraints accumulate, reflecting the growing complexity of each successive solve. Additionally, ProtLib-Designer is inherently sequential: each iteration depends on the outcomes of previous iterations, preventing parallelization. As a result, for larger libraries (e.g., >1,000 sequences), the total generation time grows substantially.

**Figure 14:**
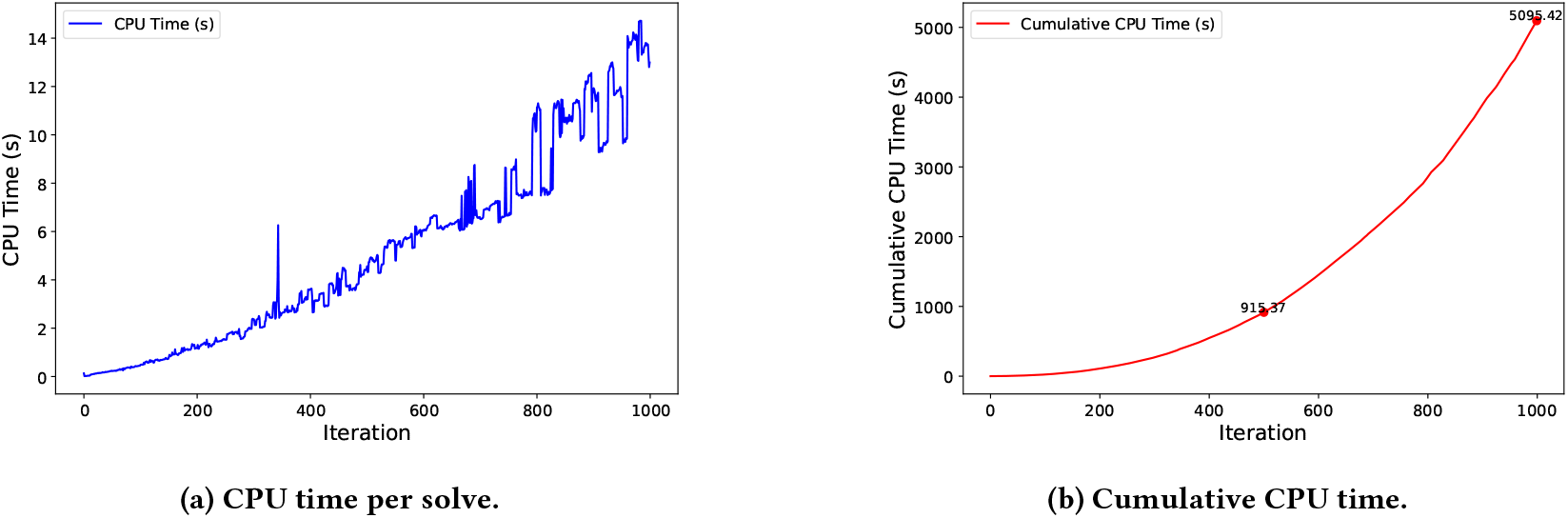
Computational cost of ProtLib-Designer. (a) CPU time in seconds for each sequential solve. (b) Cumulative CPU time across all solves.

**Figure 15:**
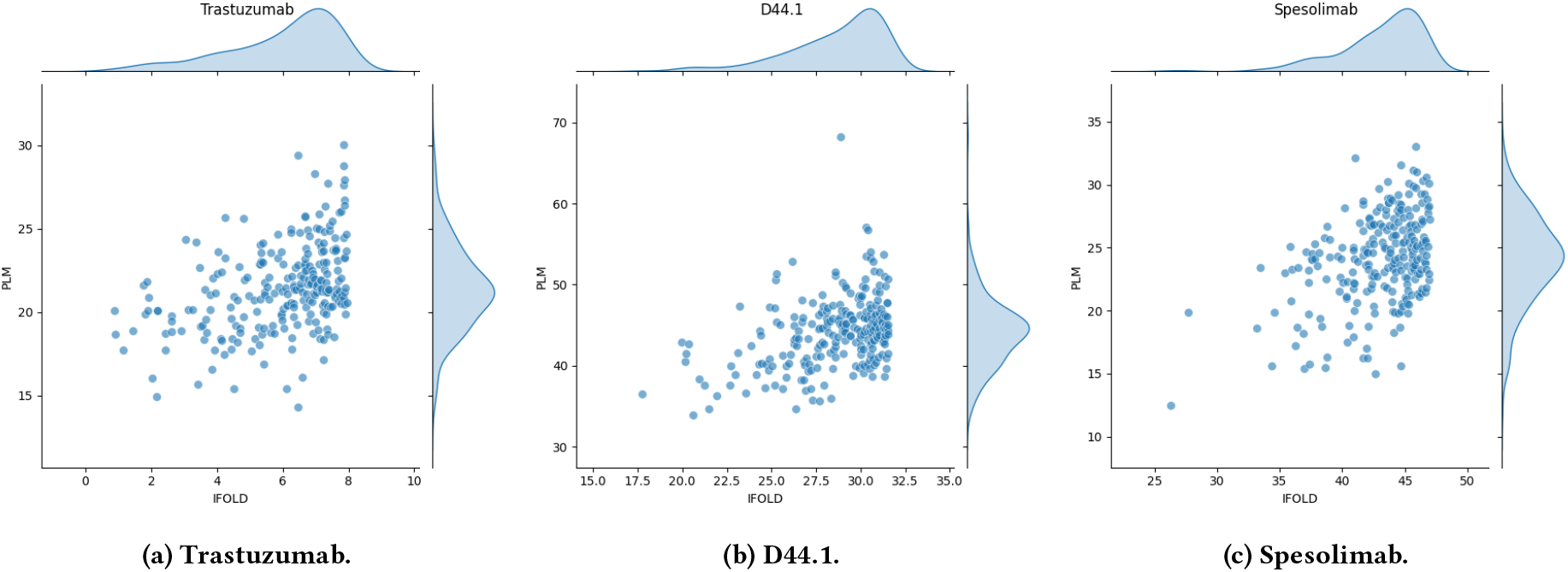
Performance of the Multi-Objective GFlowNet trained policy on each of the benchmark systems.

It is important to note that our current ILP solver uses the opensource PuLP library (https://coin-or.github.io/pulp/), which is not parallelized and is known to be less efficient than industrial-grade solvers such as Gurobi or CPLEX. These solvers can handle ILPs with thousands of constraints much more rapidly, often in seconds. We wanted to provide a full open source implementation, which is why we chose PuLP. However, these times are not representative of the performance of more advanced solvers.

#### ProtLib-Designer solve-and-remove enables computation of the Pareto front

Methods that utilize a linear scalarization are known to only be able to recover solutions that lie on the convex regions of the Pareto front, i.e the Convex hull [41]. Algorithms that use a weighted-sum scalarization are limited in this manner. However, the solve-and-remove algorithm enable ProtLib-Designer to compute solutions that lie in concave regions on the Pareto front, and in our experiments recovered the full Pareto front. The solve-and-remove algorithm alters the shape of the solution landscape ensuring that previously selected solutions are removed from consideration. Throughout the execution of the algorithm, the landscape changes, and as solutions are removed regions of the landscape becomes convex. Therefore, previously concave regions become convex allowing for ProtLib-Designer to compute the given solution using the weighted sum methods.

#### Additional Experiments with Multi-Objective GFlowNets

We adapted the multi-objective GFlowNet framework from Jain et al. [24] to our antibody-engineering context, including custom handling of mutation budgets and region-specific sampling. In our setup, the policy proceeds left-to-right through the user-defined mutable region, choosing either the wild-type or a mutant residue at each position. Sampling the wild-type amino acid “skips” that site without counting against the maximum mutation allowance. Once the budget is exhausted, further sampling terminates without altering the reward.

To integrate our ProtLib-Designer reward signal—which can be negative—into the GFlowNet’s positive-reward paradigm, we introduced a custom task that linearly rescales each model score into the (0–100) interval. These transformations ensure compatibility with the GFlowNet objective while preserving relative ranking of candidates.

After training on our custom task, we sampled 1,000 sequences from the GFlowNet policy and scored them with the same evaluation pipeline as ProtLib-Designer and the other baselines (see table 6). Comparing across Trastuzumab through Spesolimab (tables 1 to 3), the GFlowNet library falls short against all baselines. We attribute this gap to the policy’s rigid left-to-right generation, which cannot focus mutations on the most critical mutable positions. Furthermore,for D44.1 and Spesolimab, GFlowNet performs very poorly with respect to the hypervolume, receiving a score of 0. In this case, the compute Pareto front contains points out of range of the chosen reference point for the given problem evaluation. The issue stems from GFlowNets inability to sample mutations with high fitness. For D44.1 and Spesolimab benchmark problems, GFlowNets samples mutations that have positive values. Given we aim to minimize the reward functions, GFlowNets clearly struggles to optimize appropriately for the problems at hand. This is further reinforced by the high BEU scores in Table 6.

**Table 6:** Diversity and fitness metrics for the libraries generated by GFlowNet for all three systems.

| System | # of unique sequences ↑ | Residue entropy ↑ | BEU ↓ | HV ↑ | Average humanness ↑ |
| --- | --- | --- | --- | --- | --- |
| Trastuzumab | 1,000 | 4.99 | 12.42 | 1681.66 | -1779.05 |
| D44.1 | 1,000 | 5.00 | 26.69 | 0.0 | -1852.62 |
| Spesolimab | 1,000 | 5.00 | 24.11 | 0.0 | -993.27 |

## Footnotes

1 We evaluate our choice of deep learning models in the Appendix.

2 All experimental data can be found here: https://github.com/dahjan/DMS_opt

